# Endoplasmic Reticulum Geometry Dictates Neuronal Bursting via Calcium Store Refill Rates

**DOI:** 10.1101/2025.10.06.677012

**Authors:** Valentina Davi, Pierre Parutto, Cecile Crapart, Yuyi Zhang, Tasuku Konno, Joseph Chambers, John P Franklin, Daniel Maddison, Mosab Ali Awadelkareem, Michael J Devine, Elena Koslover, Edward Avezov

## Abstract

The endoplasmic reticulum (ER)’s continuous morphology is tightly controlled by ER-shaping proteins, whose genetic or expression defects drive a spectrum of neurodegenerative disorders from Hereditary Spastic Paraplegia to Alzheimer’s disease. Why perturbations in ER morphology manifest specifically in neurons remains unknown.

Here, by coupling visualisation of global sub-Hz firing bursts to ER ultrastructural manipulations in hiPSC-derived cortical neurons, alongside physical simulations, we establish a key ER structure-function principle: neuronal ER architecture dictates Ca^2+^ replenishment speed. Altering ER structure hinders network ER luminal connectivity and Ca^2+^ propagation from refill points at plasma membrane contact sites, impairing the ER’s capability to supply repetitive Ca^2+^ bursts. The ER morpho-regulatory control of Ca^2+^ refill speed thus constitutes a switch on neuronal activity. These results expose the selective vulnerability of Ca^2+^-firing cells to ER structural disruptions, rationalising ER dysfunction in neurodegeneration. This mechanism could apply universally to Ca^2+^-firing cells.

## Introduction

The endoplasmic reticulum is the only organelle that spans an entire neuron—threading soma, dendrites and axons as one contiguous network of 80-30 nm-wide tubules^1–3^. The membrane-curvature factors that sculpt this architecture — reticulons, REEPs, atlastins and spastin^4–6^ — carry pathogenic mutations that cause hereditary spastic paraplegia^7–9^ and recur as de-novo variants or mis-expressed genes in other neurodegenerative pathologies including amyotrophic lateral sclerosis^10–12^ and Alzheimer’s disease^12–14^. Although these ER-morphogens are expressed across tissues, manifestation of their disruption concentrates in neurons. This raises the question of what unique functional feature of ER architecture makes neurons so vulnerable to its perturbation.

The ER’s continuous lumen allows soluble cargos — ions, metabolites, chaperones — to move rapidly between distant cellular regions^15,16^. Among those cargos, Ca²⁺ is extraordinary: the ER maintains Ca²⁺ concentrations four orders of magnitude higher than the bordering cytoplasm - in the form of sub-millimolar free Ca²⁺, and about twentyfold equivalent bound to the protein Ca^2+^ capacitors, buffering its concentration. These endow the ER with both the depth and mobility of stores needed to operate as the cell’s dominant internal Ca^2+^ handling system^17^. Ca²⁺ is a universal second messenger, and neurons exploit it to control electrophysiological signalling^18^. We therefore postulate that the ER’s continuous tubular lattice is optimised for rapid mobility of luminal solute, and that disrupting this architecture would cripple Ca²⁺ mobilisation, disproportionately affecting cells with expansive geometry and relying on frequent Ca^2+^ transients. Thus, preserving tubular continuity would be critical for neurons because it enables swift, long-range redistribution of luminal Ca²⁺ (and other contents) to sites of demand.

Investigating the plausibility of this idea first requires determining whether and how neurons deploy ER Ca²⁺ during neurophysiological activities. Neuronal Ca²⁺ signalling spans two extremes: millisecond nanodomains beneath pre- and post-synaptic zones arise from Ca^2+^ influx via the plasma membrane^19^. ER participation in these fast, low-amplitude events is partial: store depletion can leave short-term synaptic plasticity and action potential-induced Ca^2+^ transients intact^20,21^, whereas other work reports that ER presence correlates with boosts of spine transients and synaptic strengths^22–26^. At the opposite end, seconds-long, high-amplitude Ca^2+^ waves that accompany electrophysiological bursts recur at sub-hertz frequencies during cortical development^27,28^ and in coordinate regional activities in adult brains^29–32^. In theory, a sustained influx of Ca^2+^ down the gradient across the plasma membrane alone could fuel such global events and whether the ER contributes to them is unknown. Other excitable tissues with comparable time scale and magnitude firing involve the ER: myocytes’ ER (sarcoplasmic reticulum) is indispensable for excitation–contraction in cardiac and skeletal muscle^33–35^, and the ER also supports astrocytes’ Ca²⁺ waves^36^.

Here we combine orthogonal perturbations of ER morphology with Ca²⁺ imaging in human cortical neurons, supported by an in-silico model of ER architecture and its contribution to neurophysiology. We find that the ER critically fuels sub-hertz neuronal network bursts with supply of Ca^2+^. Slowing intra-ER Ca²⁺ carrier-protein mobility via orthogonal manipulations to tubular continuity disrupts the normal firing regime. Studying the refill kinetics dependence of ER structural integrity revealed that interrupting the tubular continuity doubles the half time of Ca²⁺ store refill from plasma membrane-ER contact sites, necessary for its replenishment after a firing event. This leaves the lumen under-supplied and unable to keep up with Ca^2+^ demand during repetitive activity, thus abolishing the sub-Hz, high-amplitude network bursts. The bursts’ perturbation thus occurs even though fast local Ca^2+^ transients of synaptic activity, fuelled by the extracellular pool, can function normally and the neurons are still excitable. The principle of the ER architecture’s importance to sustaining timely Ca^2+^ resupply during firing is applicable to any other tissue using global firing for function, including astroglia and muscle (known for similar scale ER Ca^2+^ firing ^33–36^).

## Results

### ER Ca²⁺ release via IP₃ receptors fuels synchronous network bursts in human iPSC-derived cortical neurons

Synchronous network bursts at sub-Hz scale appear initially in developing cortical circuits^27,28^ and locally in adults (e.g. in visual cortex and hippocampus’ dentate gyrus)^29–32^. To establish a quantifiable read-out of this activity, we differentiated NGN2-induced human cortical neurons^37^ (*iNeurons,* Fig.1A) to a point when the cultures formed synaptically connected networks that generated culture-wide high-amplitude slow Ca²⁺ transients (mean duration - full width at half maximum = 2.5 ± 0.5 seconds, inter-burst interval = 15.8 ± 3.8 s, Fig.1B-C, Video 1). Each optical Ca^2+^ burst mirrored action-potential trains detectable by multielectrode arrays (Fig. 1B). The addition of a voltage-gated sodium channel blocker, Tetrodotoxin (TTX) or synaptic inhibitors, the AMPA/NMDA blockers CNQX + MK-801 abolished the Ca^2+^ bursts (Fig. S1A-D), confirming that the readout reflects canonical, action-potential-driven network activity. Its measurable parameters — synchrony, amplitude and frequency — enable determining the impact of specific, manipulatable, organelle biology aspects on neuronal activity.

**Figure 1:**
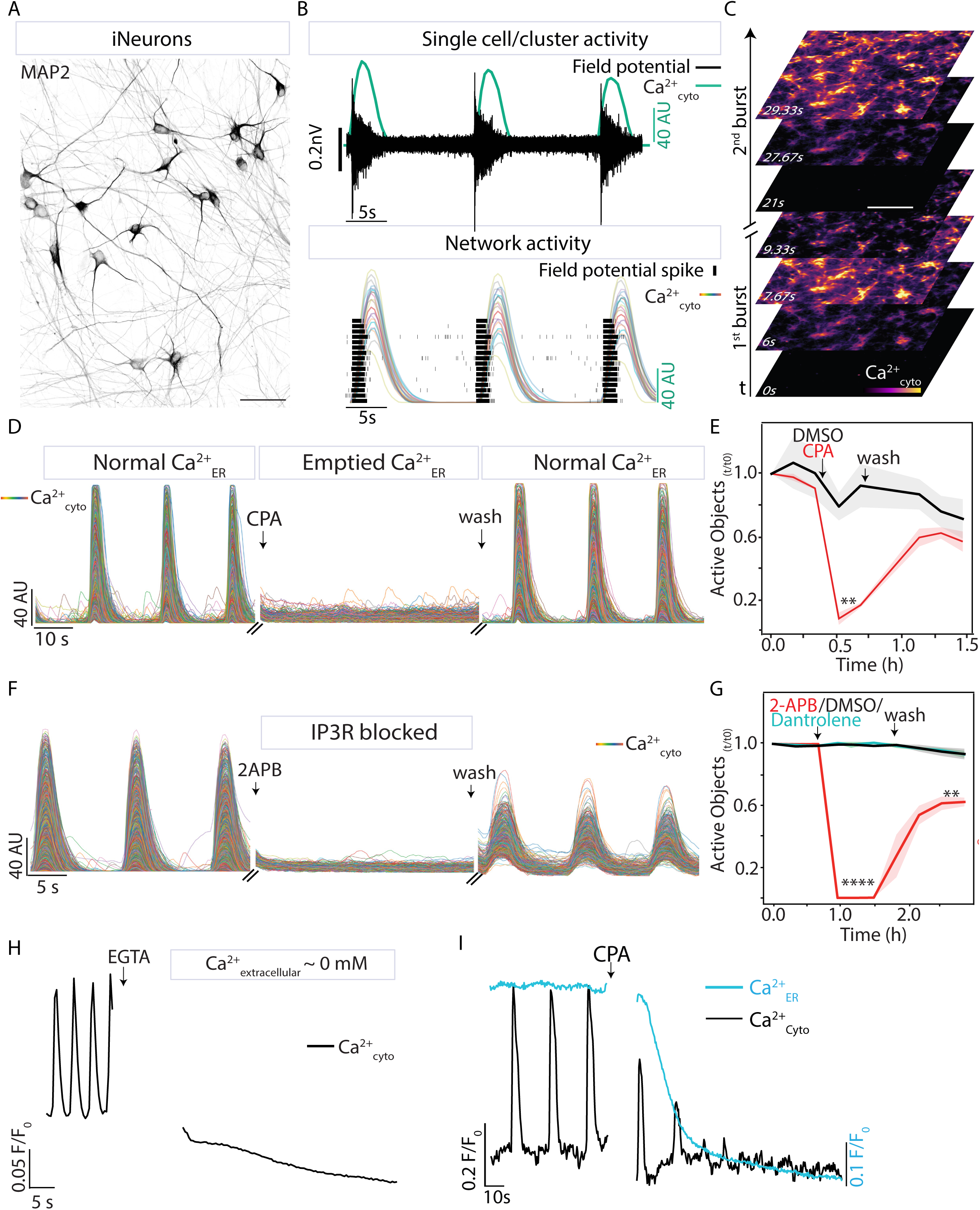
iNeurons’ network activity is fuelled by ER Ca^2+^ in an IP3R-dependent manner. **A.** Micrograph of iNeurons after 21 days of differentiation, stained with microtubule-associated protein 2 (MAP2, neuronal marker). Scale bar: 50 µm **B.** Field potential detected from a single electrode of a Multi-Electrodes Array (MEA) over time in iNeurons (top) and temporal raster plot of detected spikes from multiple electrodes (bottom), both overlayed with single iNeurons’ cytosolic Ca^2+^ traces (detected as in C). **C.** Time series of cytosolic Ca^2+^ detected by an mRuby-based Ca^2+^ sensor (Neuroburst) in iNeurons differentiated as in (A). Scale bar: 200 µm **D.** Single-cell cytosolic Ca^2+^ traces detected as in (C) from a representative culture before, 1 min after CPA treatment (SERCA pump blocker, 25 µM), and 5 min after wash-out. **E.** Quantification of firing cells (Active Objects) normalised to T_0_ over time in CPA (D) and DMSO treated cultures (solid line: mean, shade: STD, n = 3 wells). **F.** Single-cell cytosolic Ca^2+^ traces detected as in (C) treated with 2-APB (IP3R blocker, 10 µM) as in (D). **G.** Quantification of firing cells as in (E) in 2-APB (F), Dantrolene (10 µM) and DMSO-treated cultures. **H.** Cytosolic Ca^2+^ trace detected as in (C) during treatment with EGTA (3 mM). **I.** Cytosolic Ca^2+^ (black) and ER Ca^2+^ (cyan) traces simultaneously detected through Neuroburst and ER-GCaMP6_150_ respectively during CPA (25 µM) treatment. **p < 0.01, ****p<0.0001 from Student’s t-test.

Using this system, we first asked whether intracellular Ca²⁺ stores support these high-amplitude, slow (compared to the millisecond-scale local synaptic transients) and coordinated bursts. The millimolar extracellular Ca²⁺ concentration is sufficient in principle to explain the cytosolic rise. However, ER Ca^2+^ contributes to global waves in muscle and glia cells^34–36,38^. Therefore, it stands to reason that the system controlling neuronal firings on a similar spacetime scale is also designed to draw on ER-supplied Ca^2+^. Indeed, depleting the ER Ca^2+^ by irreversible inhibition of SERCA – the pump responsible for ER Ca^2+^ re-uptake from the cytosol – through Thapsigargin (TG) stopped Ca^2+^ firing (Fig. S1E, F). A reversible SERCA inhibitor, cyclopiazonic acid (CPA) caused an identical quenching of burst activity paralleled by a reduced ER Ca^2+^ concentration, but this time the bursts recovered immediately after the drug wash-out and ER Ca^2+^ replenishment. (Fig. 1D, E, I, Video 2). A Ca²⁺-loaded ER is therefore indispensable for these global network events. This conclusion is strengthened by the sensitivity of the neurons’ firing ability to blocking one of the ER Ca^2+^ release channels – IP₃-receptor – with its antagonist 2-APB. The drug eliminated Ca^2+^ firing with the same speed and completeness as SERCA inhibition (Fig. 1F, G, further confirmed in Fig. S1H-K), whereas the ryanodine-receptor (RyR) blocker Dantrolene had no effect (Fig. 1G & Fig. S1G). This result is consistent with IP3-dependency of Ca^2+^ waves in other neuronal subtypes, and the absence of reports of RyRs-mediated Ca^2+^ waves in neurons^39^.

Notably, chelating extracellular Ca²⁺ with 3 mM EGTA instantaneously terminated bursts (Fig. 1H), contrasting with a gradual run-down of burst amplitude caused by blocking the ER-specific uptake via SERCA inhibition (Fig. 1I). This is consistent with a two-step causal chain— extracellular Ca²⁺ influx triggering synaptic vesicles release, leading to activation of mGluRs and IP₃R-gated ER Ca^2+^ release to sustain high-amplitude, durable bursts. The dependence of the neuronal network activity on ER Ca^2+^ release establishes it as the quantitative baseline for testing if and how ER geometry influences neuronal behaviour.

### ER fragmentation abolishes synchronous network firing

Having documented that ER Ca^2+^ fuels global firing, we next examined whether the continuous tubular architecture of the organelle is required for this neuronal activity. We perturbed ER morphology in two orthogonal ways: first, we overexpressed the short isoform of the ER membrane protein Reticulon 3 (RTN3a), which leads to ER vesiculation while avoiding any indirect effect from its functional cytosolic tail, which acts as an ER-phagy adaptor^40^ (Fig. 2B). Second, we attained similar structural ER perturbation by orthogonal means, this time provoking it from within the ER lumen. We overexpressed the dementia-linked mutant Neuroserpin (NS_G392E_, cause of Familial encephalopathy with neuroserpin inclusion bodies - FENIB)^41,42^, to induce ER vesiculation via protein accumulation in the lumen – an ER distortion similar to that caused by its analogous serpin, α1-antitrypsin^43^ (Fig. 2C). We used these manipulations to convert the normal reticular network (Fig. 2A) into vesicular, scantily interconnected ER-clusters— as visualised by live-cell fluorescence (Fig. 2B-C) and electron microscopy of iNeurons (Fig. 2D; for assessment of luminal connectivity between clusters, see Fig. 3A-C). These manipulations left survival-vital ER functions apparently unchanged, at least within the timeframe- of subsequent experiments (Fig. S2A, B).

**Figure 2:**
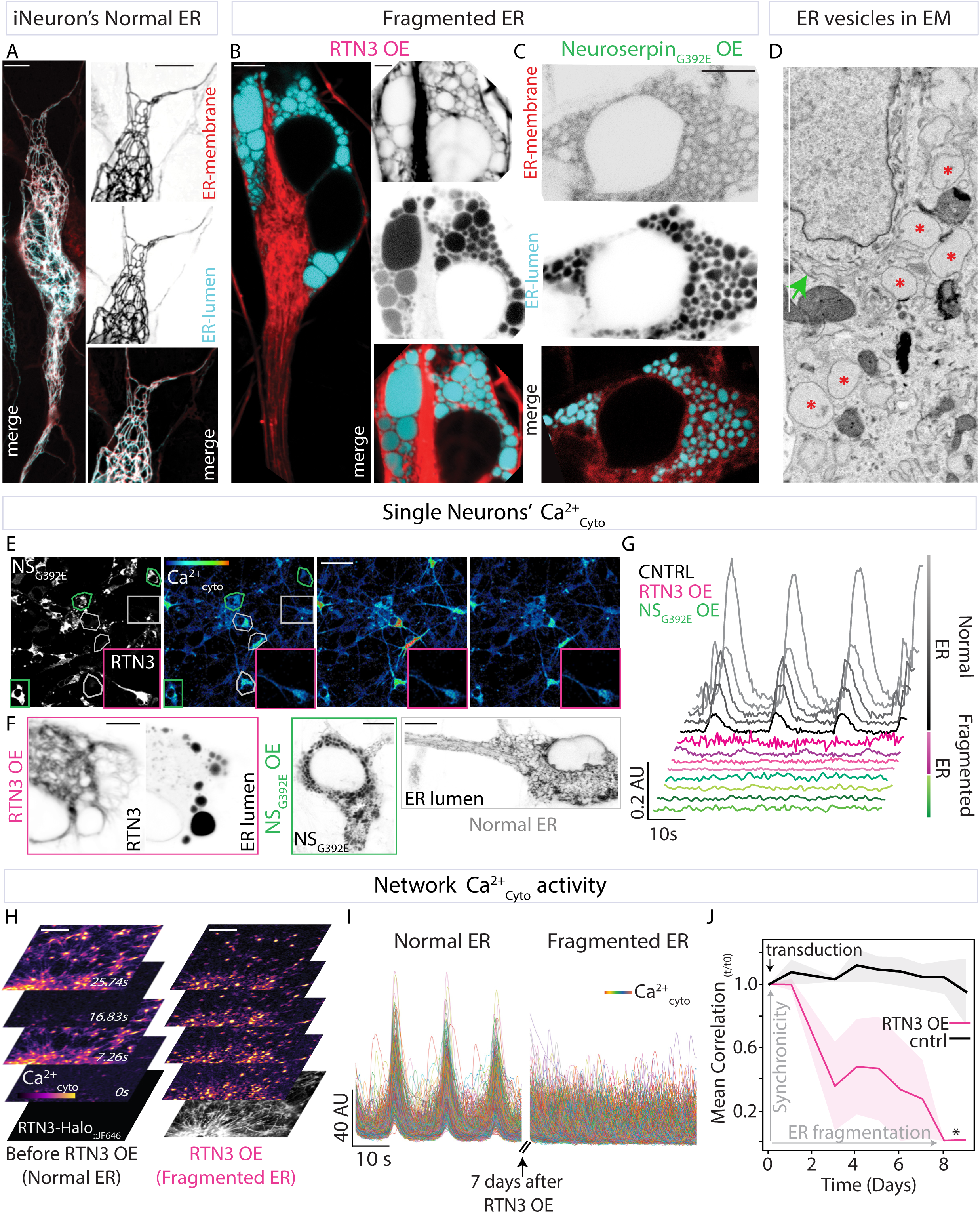
ER with perturbed morphology fails to support iNeurons’ cytosolic Ca^2+^ bursts. **A-C.** Micrographs of the ER in iNeurons with exogenously expressed **A.** ER membrane protein Halo-Sec61β labelled with JF646 (red) and ER luminal marker mEmerald-KDEL (cyan), **B.** ER membrane protein RTN3a-Halo labelled with JF646 (red) and ER luminal marker D4ER (cyan), **C.** Halo-NS_G392E_ labelled with JF646 (cyan) and ER membrane marker ER-targeted GCaMP3 (red). Note the accumulation of ER luminal content in vesicles in (B) and (C). **D.** Scanning electron micrograph of iNeurons with deformed ER as in (C) (Red stars: ER vesicles, green arrow: normal ER). Scale bars A-D: 5 µm**. E.** Micrographs of iNeurons with sparse exogenous expression of Halo-NS_G392E_ or RTN3a-Halo (inset), both labelled with JF650 (left) and time series of the Ca^2+^_cyto_ detected as in Fig.1C in the same field of view (right). Magenta Regions of Interest (ROI): RTN3OE cell, Green ROIs: NS_G392E_OE cells, Grey ROIs: controls (not infected). Scale bar: 100 µm. **F.** Magnifications of rectangular ROIs in (E). RTN3 and NS_G392E_ are labelled as in (E). ER lumen is visualised by exogenous expression of ER-mEmerald. Scale bars: 5 µm. **G.** Single-cell cytosolic Ca^2+^ traces of CNTRL (black), RTN3aOE (magenta) and NS_G392E_OE (green) iNeurons from (E). **H.** Time series of Ca^2+^_cyto_ detected as in Fig. 1C in iNeurons before (left) and after 7 days of exogenous expression of RTN3a (right). Note the loss of synchronous activity. Scale bars: 50 µm **I.** Single-cell cytosolic Ca^2+^ traces from (G). **J.** Mean time correlation between traces as in (I) over time (*p<0.05 from Student’s t-test, n = 3 wells).

**Figure 3:**
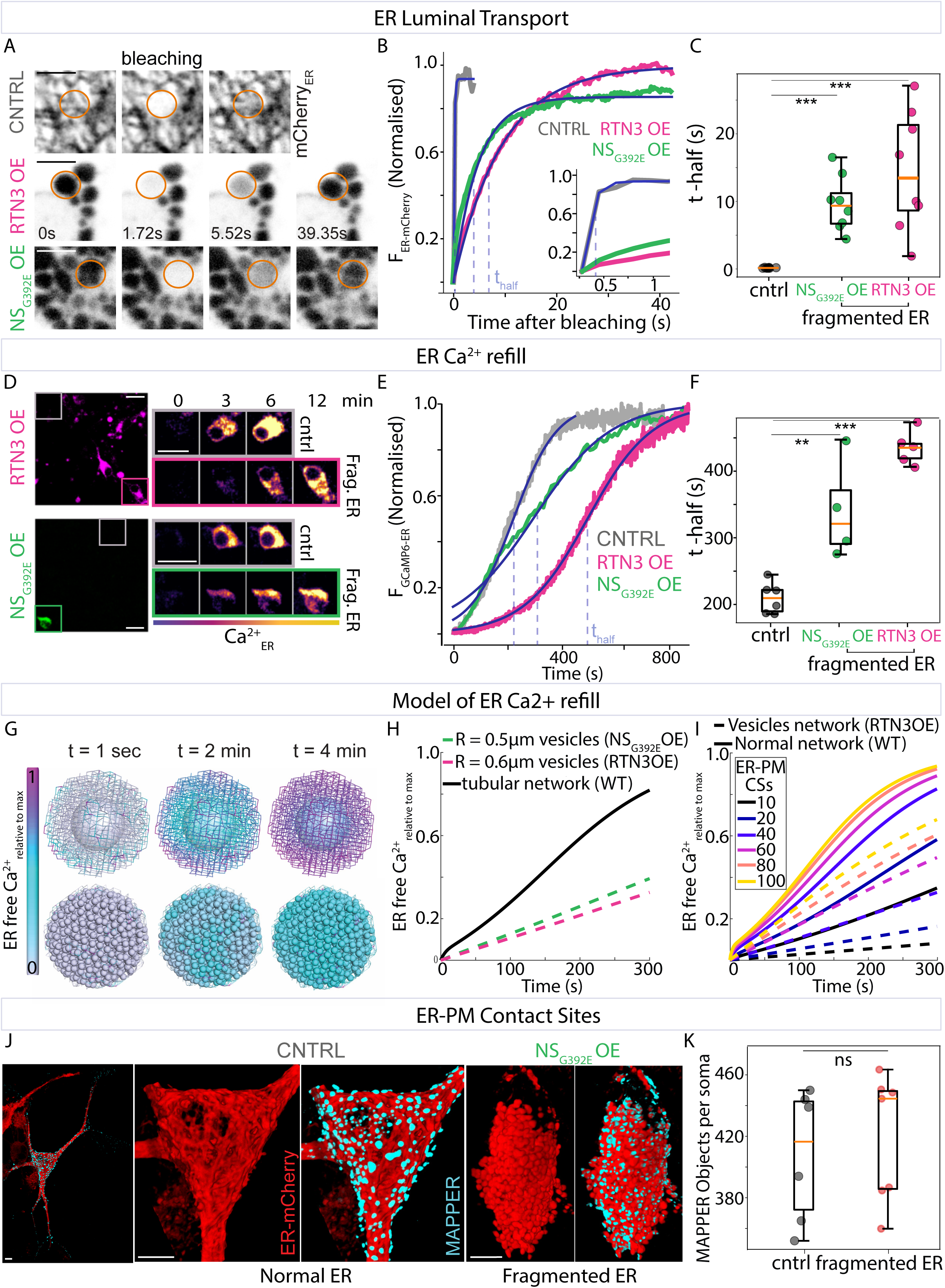
ER structure defines ER Ca^2+^ refill. **A.** Fluorescence Recovery After Photobleaching (FRAP) time series of the ER luminal protein mCherry-KDEL in normal (top) and fragmented ER (RTN3aOE middle, NS_G392E_OE bottom) in iNeurons. The orange circles represent the bleached area. Scale bars: 2 µm. **B.** mCherry-KDEL intensity traces after photobleaching as in (A) (black: control (not infected), magenta: RTN3aOE, green: NS_G392E_OE). Blue curve: exponential fit, light blue dashed line: half recovery time (t-half). Inset: blow-up of the same plot. **C.** Half recovery time (t-half) values from the exponential fit as in (B) from WT (n = 8 cells), RTN3 OE (n = 8 cells, ***p = 4.43 x 10^-4^), and NS_G392E_OE (n = 8 cells, ***p = 1.16 x 10^-5^) cells. Each dot represents the t-half from a single cell. **D.** Micrographs of iNeurons with sparse exogenous expression of RTN3a-Halo (top, magenta) and Halo-NS_G392E_ (bottom, green), both labelled with JF646 (left) and time series of the ER Ca^2+^ sensor GCaMP6_150_ in the same field of view (right) after washout of BTP2 (ORAI1 blocker, 10 µM, 20 min). Magenta ROI: RTN3aOE cell, Green ROI: NS_G392E_OE cell, Grey ROIs: controls (not infected). Scale bars: 20 µm. **E.** Single-cell ER Ca^2+^ trace detected and treated as in (D) of control (black), RTN3aOE (magenta) and NS_G392E_OE (green) iNeurons. Blue curve: sigmoid fit, light blue dashed line: time to half-recovery (t-half). **F.** T-half values from the sigmoid fit as in (E) of control (n = 6 cells), RTN3aOE (n = 5 cells, ***p = 1.48 x 10^-7^) and NS_G392E_OE (n = 4 cells, **p = 5.49 x 10^-3^) iNeurons. Each dot represents the t-half from a single cell. **G**. Simulation snapshots of Ca^2+^ refill from ER-PM contact sites. Colour corresponds to free luminal Ca^2+^ levels, at increasing times after refill is initiated. Top: tubular network, bottom: partially fragmented network of spherical vesicles of 0.6 µm radius connected by narrow tubes. **H**. Average free luminal Ca^2+^ throughout the network, in the spatial transport model, plotted against simulation time. Black solid line: tubular network. Dashed lines: networks of spherical vesicles with radius 0.6 µm (red) and 0.5 µm (pink), representing RTN3aOE and NS_G392E_OE ER structures, respectively. **I**. Average luminal free Ca^2+^ plotted over time for simulations with the tubular network (solid lines) and spherical vesicle network (dashed lines). Colours correspond to different number of ER-PM contact sites with fixed unbound Ca^2+^ concentration **J.** 3D reconstructions from confocal z-stacks of mCherry-KDEL (red) and MAPPER (cyan) in WT (left) and NS_G392E_OE (right) iNeurons. Scale bars: 5 µm. **K.** MAPPER clusters per cell detected as in (J) in iNeurons with normal (n = 6) and fragmented ER (NS_G392E_OE and RTN3aOE, n = 6 cells, ^ns^p = 0.9). Each dot represents MAPPER clusters from a single cell. Statistical significance was determined by Student’s t-test.

Strikingly, within the synchronously firing networks, iNeurons with ER fragmentation induced by either misshaping factor were not active (Fig. 2E-G, Videos 3 and 4). In contrast, neighbouring cells that did not express the transgenes in the same field retained normal Ca^2+^ firing (Fig. 2E-G). At the culture level, the mean correlation coefficient—a proxy for network synchrony—declined steadily after RTN3a overexpression in the majority of cells (Fig. 2H-J, Video 5). Since the same burst-cancelling effect was attainable through two independent manipulations, similar only in their capacity to perturb ER morphology, it is reasonable to rule out the possibility that their other functions are responsible for this effect. Thus, we conclude that the loss of ER continuity prevents single-neuron Ca^2+^ firing and abolishes synchronous network firing.

### ER fragmentation hinders intra-ER Ca²⁺ transport, slowing store refill

Next, we investigated the mechanism underlying the absence of spontaneous cytosolic bursts in iNeurons with altered ER morphology. This phenotype is not explainable by defects in the ER’s ability to store or release Ca^2+^, since ER free and total Ca^2+^ remain within the normal range upon ER-fragmentation in iNeurons, as previously observed in COS-7 cells^15^ (free ER Ca^2+^ concentration in absolute terms was measured by calibrated fluorescence life time imaging, FLIM of D4ER-Tq probe (Fig. S2C-E), while total was assessed by measuring the Tg-releasable pool, (Fig. S2F, G)). Further, the amplitude and kinetics of IP₃-evoked Ca²⁺ transients in ER-perturbed neurons were indistinguishable from wild-type (Fig. S2H, I).

This leaves the kinetic consideration of Ca^2+^ delivery to ER stores as the prime parameter suspected to be dependent on ER structural integrity. To support repeated releases, the ER Ca^2+^ pool replenishment should proceed at a speed matching demand. The ER cannot be replenished solely by SERCA-mediated re-uptake of Ca^2+^ back from the cytoplasm, as the ion is also rapidly cleared to the cell exterior by plasma membrane pumps. This is evident in classical experiments showing fast decay of cytoplasmic Ca^2+^ upon SERCA inhibition (e.g.^44–46^). The critical contribution of the extracellular Ca^2+^ for store refill is further evident from the inability of ER to replenish post-release when Ca^2+^ in the extracellular medium is chelated^47^(Fig. S3).

Thus, we postulate that the ER refill rate depends on Ca^2+^ propagation from its intake points at ER-PM contact sites — hubs of store-operated Ca^2+^ entry (SOCE) from the extracellular space — to the bulk of the ER network volume. Since Ca^2+^ propagation inside the ER is partly dictated by the mobility of its high-capacity carriers^15^, we sought to assess whether the extent of reduction in protein mobility upon ER fragmentation could explain the retarded ER refill. Fluorescence Recovery After Photobleaching (FRAP) measurements showed a ∼ 10-fold reduction in luminal protein mobility (Fig. 3A-C, S4A-B), on par with luminal transport reduction observed in ER-perturbed COS-7 cells^15^. Crucially, the slower protein mobility correlated with the reduction in ER-refill rate, as visualised by the speed of Ca^2+^ recovery after its depletion caused by blocking of ER-plasma membrane refill points with an ORAI1 inhibitor (BTP-2): The wild-type neurons’ ER Ca²⁺ refill rate of t½ = 229.5 ± 34.6 s was slowed to 434.5 ± 21.6 s by RTN3a overexpression and to 341.1 ± 66.5 s by NS G392E (Fig. 3D-F, Video 6).

To correlate this slowdown in Ca^2+^ refill kinetics with perturbed ER morphology, we established an *in-silico* physical model representing transport of Ca^2+^ through the ER network. As illustrated in Fig. 3G, the wildtype ER was represented as a lattice-like network of tubules, connected to the extracellular environment at PM contact sites and to a perinuclear reservoir. The fragmented ER structures resulting from RTN3 and NS_G392E_ overexpression were represented as a network of larger and smaller spherical vesicles, respectively, connected to nearest neighbours by narrow short tubes. The vesicles serve as traps for diffusive particles, which must find narrow tubule entrances to escape. Notably, estimates of diffusive transport imply that the effective rate of spread through the network of spheres is expected to be significantly reduced compared to a tubular network (see Modelling in Methods for detail), and that larger vesicles should yield slower transport, supporting our experimental findings (Fig. 3A-C).

We employed numerical simulations to explicitly compute refill rates for these distinct network structures, assuming a fixed free Ca^2+^ concentration at PM contact sites and an initially empty ER lumen. The tubular structure was found to fill substantially faster than the partially fragmented structures, with larger vesicles showing slower refill (Fig.3G-H).

These refill rates are limited by a combination of factors, including bulk transport through the 3D network (substantially slower in the vesicular structures) and the number of contacts with the plasma membrane (Fig. 3I). To rule out the possibility that the impaired refill in neurons with fragmented ER was caused by disruption of ER-PM contacts, we examined their density (visualised with MAPPER^48^) and found no change in neurons with perturbed ER (Fig. 3J, K). Since these contacts are hubs for STIM-ORAI interaction (the actuators of SOCE), this result indicates that the refill defect lies in Ca²⁺ transport within the fragmented ER rather than in SOCE engagement at the periphery. This is consistent with a previous report of contact sites persisting despite vesiculation of the ER structure^49^.

Together, the *in-silico* model and the experimental measurements of ER transport and refill show that a continuous ER network is required to move Ca²⁺ rapidly from peripheral entry points to the bulk reservoir. Thus, ER shape dictates its store refill rate.

### Slow ER refill hinders ability to generate persistent bursts

Since ER fragmentation affects Ca^2+^ refill rates, we hypothesised that cells with disrupted ER shape cannot replenish ER Ca²⁺ stores quickly enough to support repetitive global releases. To validate the plausibility of this hypothesis, we first interrogated the extent to which ER refilling limits repetitive firing. To mimic the refill bottleneck without altering ER shape, we again acutely blocked SOCE using the ORAI1 antagonist BTP2. Immediately after treatment, cytosolic Ca^2+^ bursts persisted but their amplitude gradually declined until they were completely abolished (Fig. 4A, Video 7). The progressive rundown mirrors the burst decline observed when the ER was slowly depleted with SERCA inhibitors (Fig. 1I). These are consistent with the critical contribution of ER Ca^2+^ to fuelling the bursts rather than merely triggering them. Therefore, hindering ER refill alone is sufficient to exhaust the capability of ER to sustain the Ca^2+^ firing activity. Further, repeated, high-frequency electrical field stimulation of iNeurons with RTN3a-induced ER fragmentation led to progressively declining cytosolic Ca^2+^ bursts amplitude (Fig. 4B, C) — whereas cells with normal ER from the same field of view could sustain repeated Ca^2+^ bursts of consistently high amplitude. The gradual dampening of burst amplitude mirrors the refill inhibition phenotype achieved by BTP2 — while these cells retain normal Ca^2+^ release kinetics (Fig. S2H, I). Therefore, we conclude that the slow refilling of fragmented ER causes its failure to sustain the global mode of Ca^2+^ firing.

**Figure 4:**
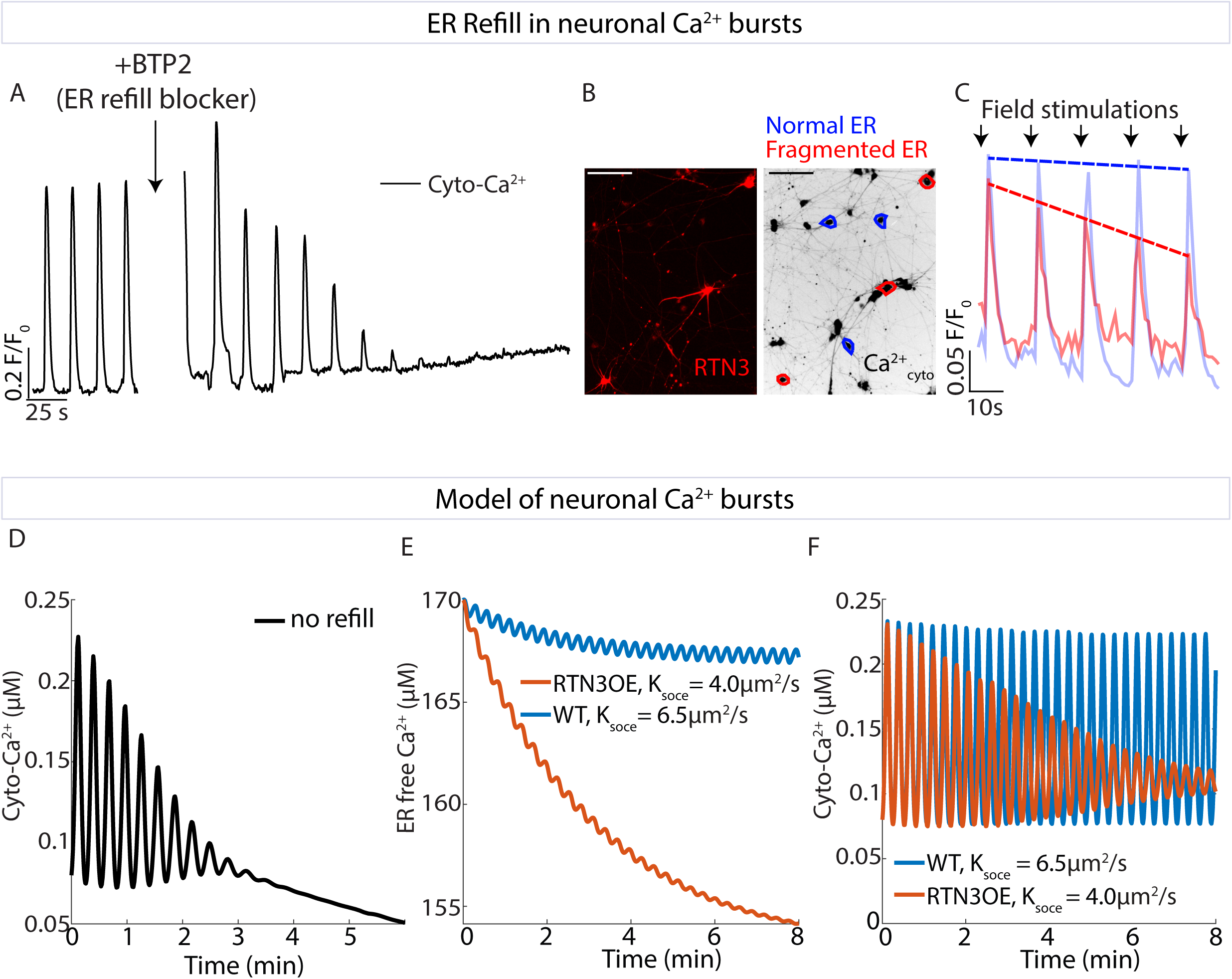
Perturbing ER refill hinders ability to generate persistent bursts. **A.** Single-cell cytosolic Ca^2+^ trace before and after treatment with BTP2 (10 µM, ORAI1 inhibitor). **B.** Micrographs of iNeurons with sparse exogenous expression of RTN3a-Halo labelled with JF646 (left) and stained with the cytosolic Ca^2+^ dye Cal-520 (right). Blue ROIs: controls (not infected), red ROIS: RTN3aOE cells. Scalebar: 100 µm. **C.** Single-cell cytosolic Ca^2+^ traces of WT (blue) and RTN3aOE (red) iNeurons detected as in (B) after periodic field stimulations (arrows, every 15s). Dotted line represents linear fit of the bursts’ peaks. **D.** Spontaneous Ca^2+^ firing in nonlinear dynamic model, with no ER refill (k_soce_ = 0), showing gradual decay of burst magnitude and eventual cessation of release. **E**. Unbound Ca^2+^ in the ER lumen, from the spontaneous Ca^2+^ firing model with two different rates of SOCE refill, as estimated from spatial simulations, representing a cell with WT (blue) versus partially fragmented (red) ER. **F**. Corresponding cytoplasmic Ca^2+^ levels in the model, showing sustained oscillations with high refill (blue) and gradual dampening with a low refill rate (red).

Our physical model quantitatively determined to what extent limited ER refill rates can account for the observed bursts’ gradual rundown. We implemented a modified version of an archetypal model for Ca^2+^ oscillations^50^, incorporating IP3-receptor mediated Ca^2+^ release from the ER, SERCA-mediated Ca^2+^ reuptake, cytoplasmic clearance, and refill of the ER via SOCE. This simplified model treats the ER and cytoplasmic Ca^2+^ pools as well mixed, without representing spatial transport effects within each compartment. It uses a Hodgkin-Huxley-like formalism to describe the time-dependent activation and inactivation of IP3R in response to cytoplasmic Ca^2+^ levels, enabling the emergence of autonomous Ca^2+^ release pulses with a periodicity of 16 sec, similar to experimental observations (for model and its parameterisation details see Modelling in Methods).

A key parameter in the model is the rate of ER refill through SOCE. If this refill process is completely shut off, then the modelled Ca^2+^ bursts gradually dampen (Fig. 4D), as observed upon BTP2 addition (Fig.4A). To represent the overall SOCE refill rate, we use an effective rate constant (k_soce_) extracted from our simulations of intra-luminal Ca^2+^ transport (Fig. S5), which decreases by approximately 40% upon ER fragmentation. This decrease results in a relatively small (<10 %) change to the ER luminal Ca^2+^ levels in the model (Fig. 4E), potentially explaining the absence of a detectable ER Ca^2+^ drop in our experiments. However, due to the nonlinear nature of the model, this small change can be sufficient to dampen out Ca^2+^ release pulses over a multi-minute timescale (Fig. 4F), matching our experimental observations (Fig 4C). While not quantitatively predictive due to the presence of multiple under-specified parameters, the Ca^2+^ pulsing model illustrates the plausibility of Ca^2+^ bursts declining due to a moderate decrease in the SOCE refill rate alone.

Collectively, pharmacological inhibitions, spontaneous and stimulated firing assays, ER Ca^2+^ dynamics measurements and mathematical simulations converge on a single conclusion: a continuous, fast-tunnelling ER network is required in neurons to replenish its Ca^2+^ during global releases; when refilling cannot keep pace with demand because of perturbed ER shape, burst amplitudes gradually decrease until individual activity ceases, thus disrupting the network firing. The high Ca^2+^ demand global sub-Hz firing, are particularly sensitive to diminishing in supply, even when the ms-scale synaptic transients sourced from extracellular Ca^2+^ pool are unaffected.

## Discussion

Our results identify a mechanism of ER Ca^2+^ refill-limited excitability**—**an architectural checkpoint for firing. This constitutes a kinetic bottleneck that caps neuronal network activity: the time required for Ca²⁺ to traverse the continuous ER lumen. Perturbing the integrity of tubular ER slows intra-luminal Ca^2+^ mobility, particularly its tunnelling from the ER plasma membrane contact sites to replenish the store. Although impeded refill is not detrimental for normal Ca^2+^ store maintenance in non-firing cells (as membrane machinery of Ca^2+^ pumps and channels remains functional in fragmented ER^15^), it is strongly felt when ER is required to support repetitive transient elevations of cytoplasmic Ca^2+^ while firing. During such global release activities, the Ca^2+^-starved ER struggles to keep up with Ca^2+^ demand and when its contribution fades away, firing stops. Thus, organelle geometry stands alongside ion channels and synaptic receptors as a first-order determinant of periodic firing, particularly at high amplitude with sub-Hz frequency.

Modelling, constrained by fluorescence-recovery measurements, shows burst probability collapsing once the refill rate slows beyond a threshold. Crucially, we observe a progressive decline in burst amplitude before complete failure: partial fragmentation or mild SOCE-mediated refill inhibition produces mis-scaled events rather than an abrupt switch-off. These suggest a possible regime of controlled “luminal starvation” acting as a physiological rheostat: neurons might transiently fragment the ER to dampen network output. Such ER transient fragmentations have been documented in (patho)physiological circumstances with attenuated neuronal activity, particularly in ischemia^51^ and after prolonged sensory overstimulation^52^. Further, corticospinal neurons combine millimetre-scale arbours, distant from the large somatic storage ER reservoirs, with an unfavourable volume-to-store ratio, making them sensitive to Ca^2+^ delivery delays. This effect can rationalise why ubiquitous ER-shaping mutations cripple neurons first and why symptoms often manifest as distal axonopathies. This is consistent with recent observations that hereditary spastic paraplegia (HSP)-mimicking loss of reticulon in flies reduce terminal Ca²⁺ transients and neurotransmitter-release probability^53,54^. The effect we described for the seconds-scale bursts can hold true for shorter time scales and local firing events in a more subtle form. Thus, instead of the all-or-nothing scenario for bursts, the effect may translate into weaker synaptic strength or reduced likelihood to fire.

Further, complete loss of ATL, another ER morphogen with loss-of-function mutations in HSP, also showed retarded refill (Fig. S6A-C).

Cardiomyocytes and skeletal muscle fibres depend on sarcoplasmic reticulum for force generation, and astrocytes rely on ER stores for their function state-associated intercellular Ca²⁺ waves^33–36^. Whether morphological fragmentation modulates contractility or glial signalling in vivo remains to be tested, but the refill speed threshold concept—and the newly recognised graded pre-collapse window—offers clear, quantitative predictions for those systems.

By establishing a direct structure–function relationship for the neuronal ER, we elevate organelle architecture to a quantitative variable in neurophysiology. The principle of refill-limited excitability suggests that modest geometric tuning—whether pathological or adaptive—can shift networks along a continuum from normal bursting to quiescence. Maintaining ER continuity therefore represents a general strategy for balancing neuronal functionality.

---

**Figure S1:**
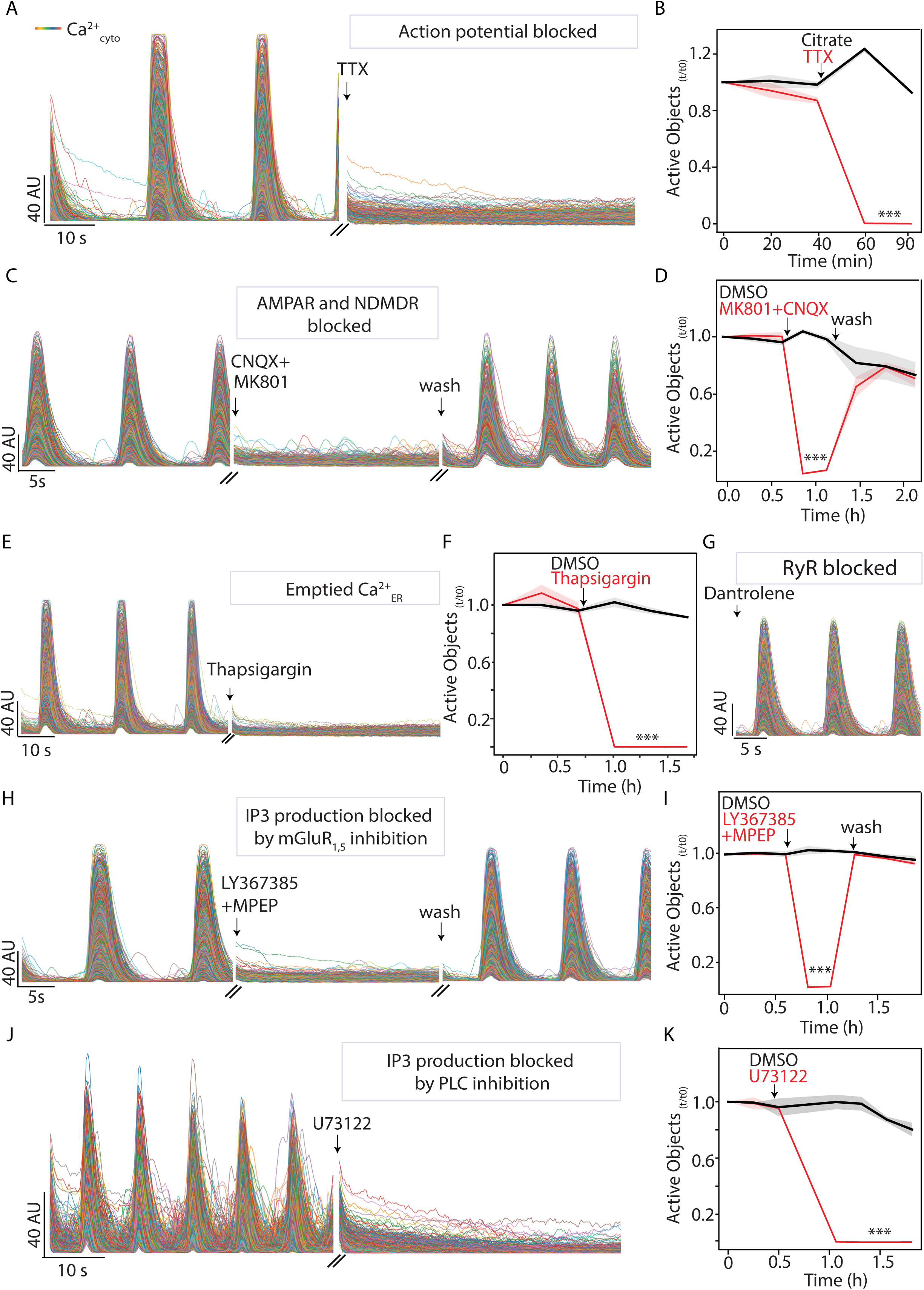
Characterisation of iNeurons’ Ca^2+^ bursts. **A.** Single-cell cytosolic Ca^2+^ traces before and 1 min after TTX treatment (1 µM). **B.** Active cells (Active Objects) count over time in TTX and Citrate treated cultures. **C.** Single-cell cytosolic Ca^2+^ traces before, 1 min after CNQX and MK801 treatment (10 µM) and 5 min after washout. **D.** Quantification of (C) as in (B). **E.** Single-cell cytosolic Ca^2+^ traces treated with Thapsigargin (2 µM) as in (B). **F.** Quantification of (E) as in (B). **G.** Single-cell cytosolic Ca^2+^ traces 5 min after Dantrolene (10 µM). **H.** Single-cell cytosolic Ca^2+^ traces treated with LY 367385 (100 µM) and MPEP (10 µM) as in (C). **I.** Quantification of (H) as in (B). **J.** Single-cell cytosolic Ca^2+^ traces treated with U73122 (PLC inhibitor, 10 µM) as in (A). **K.** Quantification of (J) as in (B). In all the experiments in this figure, iNeurons were at Day 28 and cyto-Ca^2+^ was detected through Neuroburst. Statistical significance was determined by Student’s t test; *p<0.05, **p < 0.01, ***p<0.001, ****p<0.0001. Data are represented as mean (solid line) ± STD (shade) over n = 3 wells.

**Figure S2:**
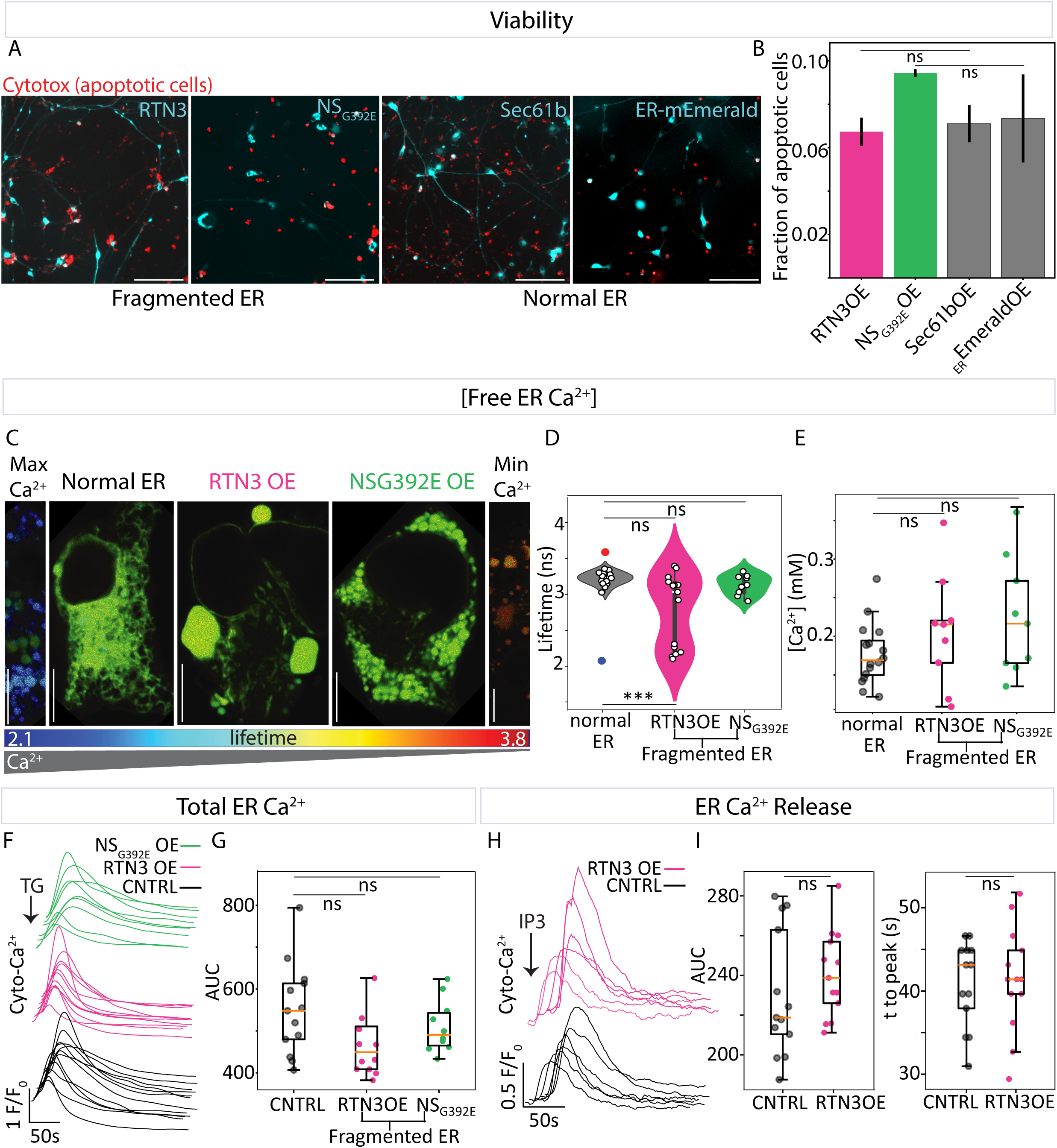
Characterisation of ER fragmentation in iNeurons. **A.** Micrographs of the apoptotic marker Incucyte^®^ Cytotox Red Dye in iNeurons after 7 days of exogenous expression of (from left to right) RTN3a-Halo, Halo-NS_G392E_, Halo-Sec61b (ER-membrane control) and mEmerald-KDEL (ER-lumen control). Halo was labelled with HaloTag Oregon Green Ligand. Scalebar: 100 μm. **B.** Fraction of cells with exogenous expressions as in (A) co-localising with the apoptotic marker (n = 3 wells, ^ns^p_RTN3_ = 0.65, ^ns^p_NSG392_ = 0.22). **C-E.** Measurements of free ER Ca^2+^ concentration. **C.** Fluorescence lifetime imaging microscopy (FLIM) images of ER-targeted D4ER-Tq Ca^2+^ FRET probe in iNeurons. From left to right: maximal ER Ca^2+^ obtained by ionomycin treatment (10 μM), WT, RTN3aOE, NS_G392E_OE, minimal ER Ca^2+^ obtained by Thapsigargin treatment (3 μM). Scalebars: 5 µm. **D.** FLIM average lifetimes detected as in (C) of WT (n = 16 cells), RTN3aOE (n = 15 cells, ^ns^p_top_ = 0.28, ***p_bottom_ = 1.33 x 10^-16^) and NS_G392E_OE (n = 9 cells, ^ns^p = 0.06) iNeurons. Each dot represents the lifetime from a single cell. Lifetimes in maximal and minimal ER Ca^2+^ conditions are indicated as red and blue dots respectively. **E.** Luminal free [Ca^2+^]_ER_ calculated from values in (D) of WT (n = 16 cells), RTN3aOE (n = 9 cells, ^ns^p = 0.20) and NS_G392E_OE (n = 9 cells, ^ns^p = 0.053) iNeurons. Each dot represents the [Ca^2+^]_ER_ from a single cell. The low-lifetime population of RTN3OE iNeurons was excluded as an artifact was suspected in this subpopulation of cells presenting extra-large vesicles. **F-G.** Measurement of Total ER Ca^2+^ load. **F.** Single-cell cytoplasmic Ca^2+^ traces detected by Cal-520 after ER release by 3 μM Thapsigargin in WT (black), RTN3 OE (pink) and NS_G392E_OE (green) iNeurons. **G**. Integrated fluorescence intensity (Area Under the Curve – AUC) of traces in (F) of WT (n = 13 cells), RTN3 OE (n = 12 cells, ^ns^p = 0.22) and NS_G392E_OE (n = 10 cells, ^ns^p = 0.31) iNeurons. Each dot represents the AUC from a single cell. **H.** Single-cell cytoplasmic Ca^2+^ traces detected by GCaMP8 after ER release through light-induced IP3 uncaging in whole field of view containing WT (black) and RTN3 OE (magenta) iNeurons. **I.** Integrated fluorescence intensity (AUC, left, ^ns^p=0.35) and time to peak (right, ^ns^p=0.82) of WT (n = 13 cells) and RTN3 OE (n = 13 cells) iNeurons. Each dot represents AUC and time-to-peak from a single cell. Statistical significance was determined Student’s t test.

**Figure S3:**
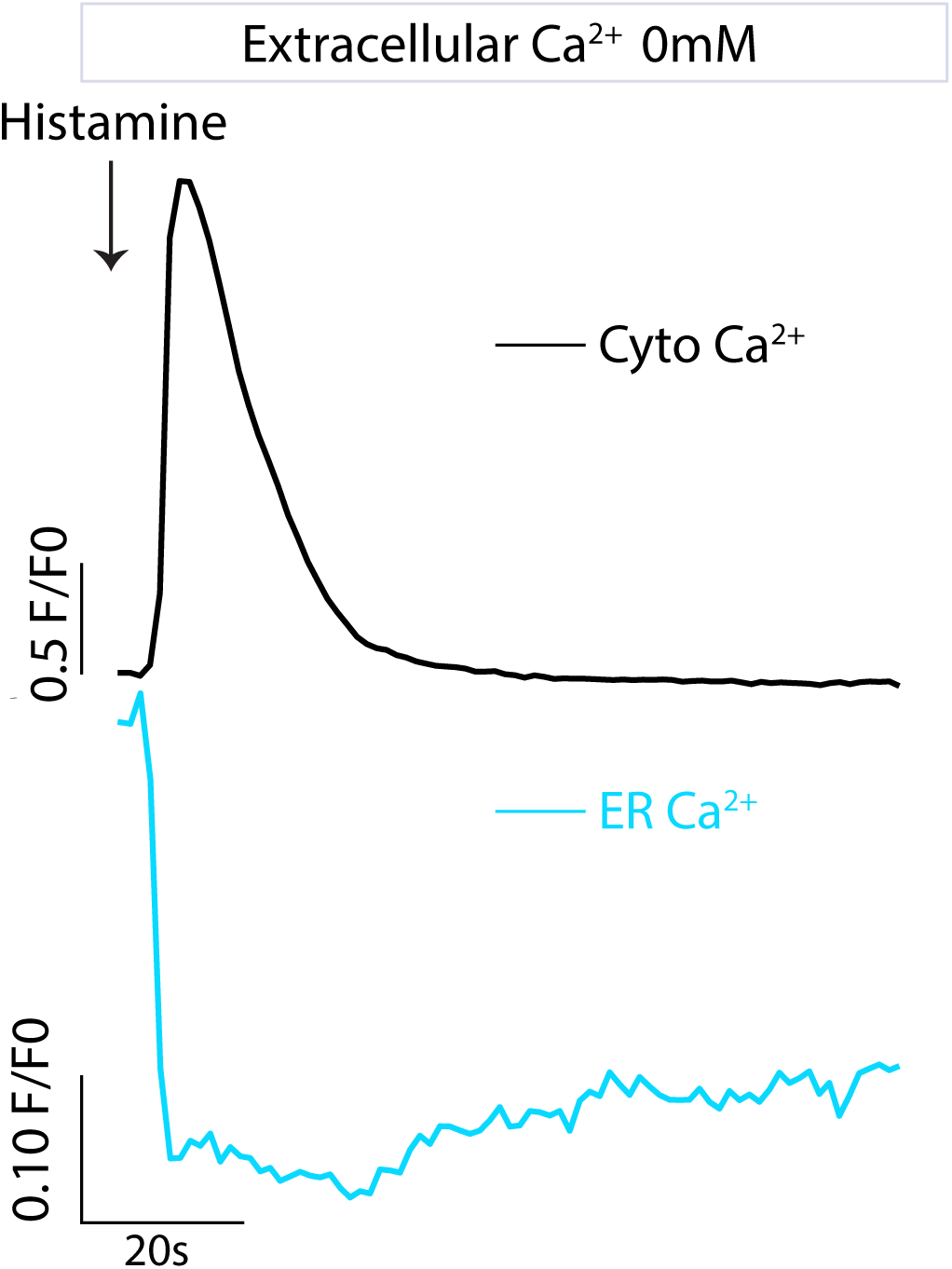
ER does not re-uptake cytosolic Ca^2+^. Single-cell cytosolic Ca^2+^ (black) and ER Ca^2+^ (cyan) traces simultaneously detected in COS7 cells through GCaMP8 and R-CEPIA_ER_ respectively. Arrow represents time of treatment with Histamine (100 µM). EGTA (3 mM) was added to the media 5 mins before the experiment.

**Figure S4:**
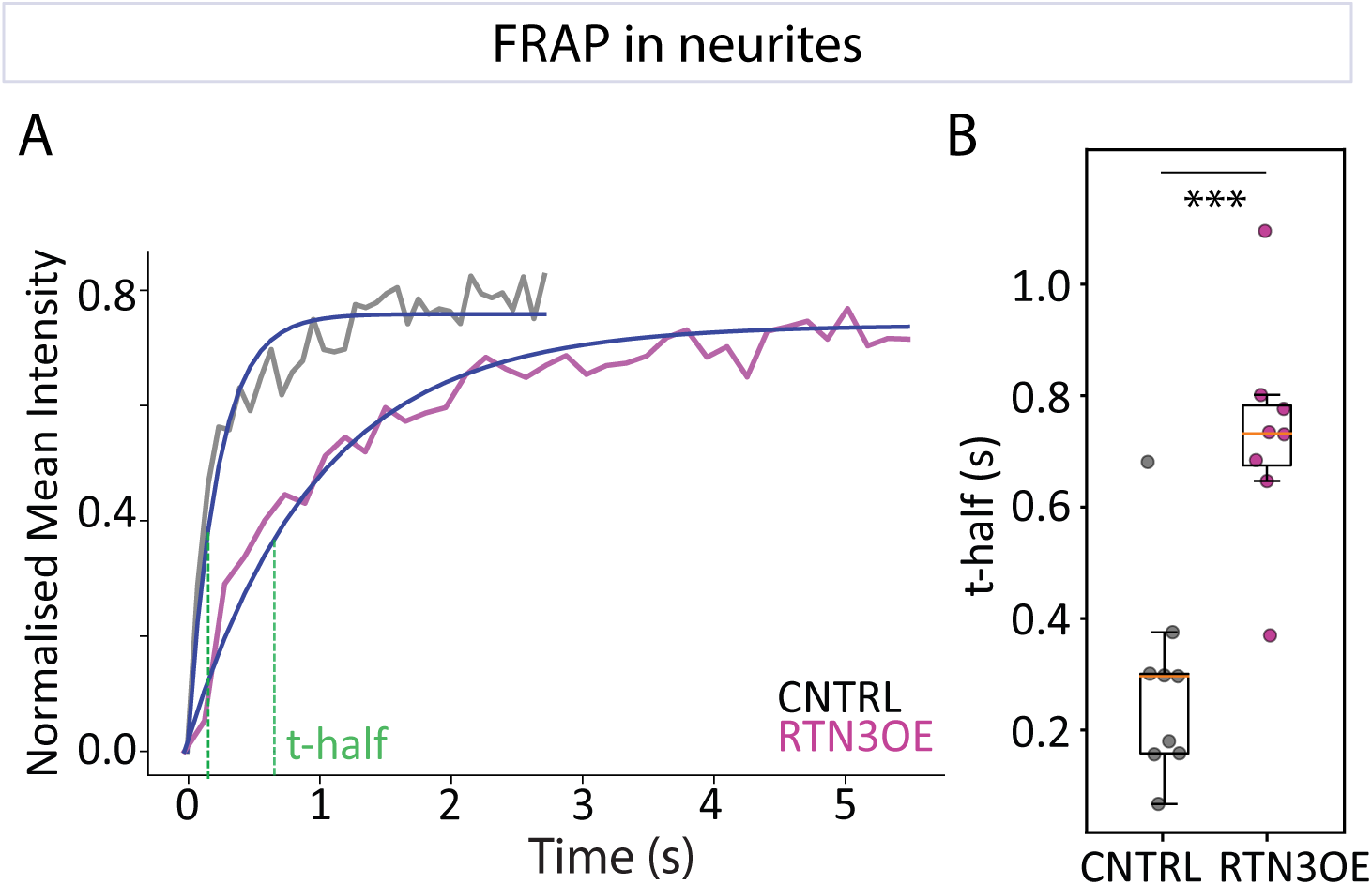
ER fragmentation affects luminal connectivity in neurites. **A.** ER luminal protein mEmerald-KDEL intensity traces after photobleaching (grey: control (not infected), magenta: RTN3aOE). Blue curve: exponential fit, green dashed line: half recovery time (t-half). **B.** Half recovery time (t-half) values from the exponential fit as in (A) from WT (n = 9 cells) and RTN3 OE (n = 8 cells, ***p = 1.9 x 10^-4^). Each dot represents the t-half from a single cell.

**Figure S5:**
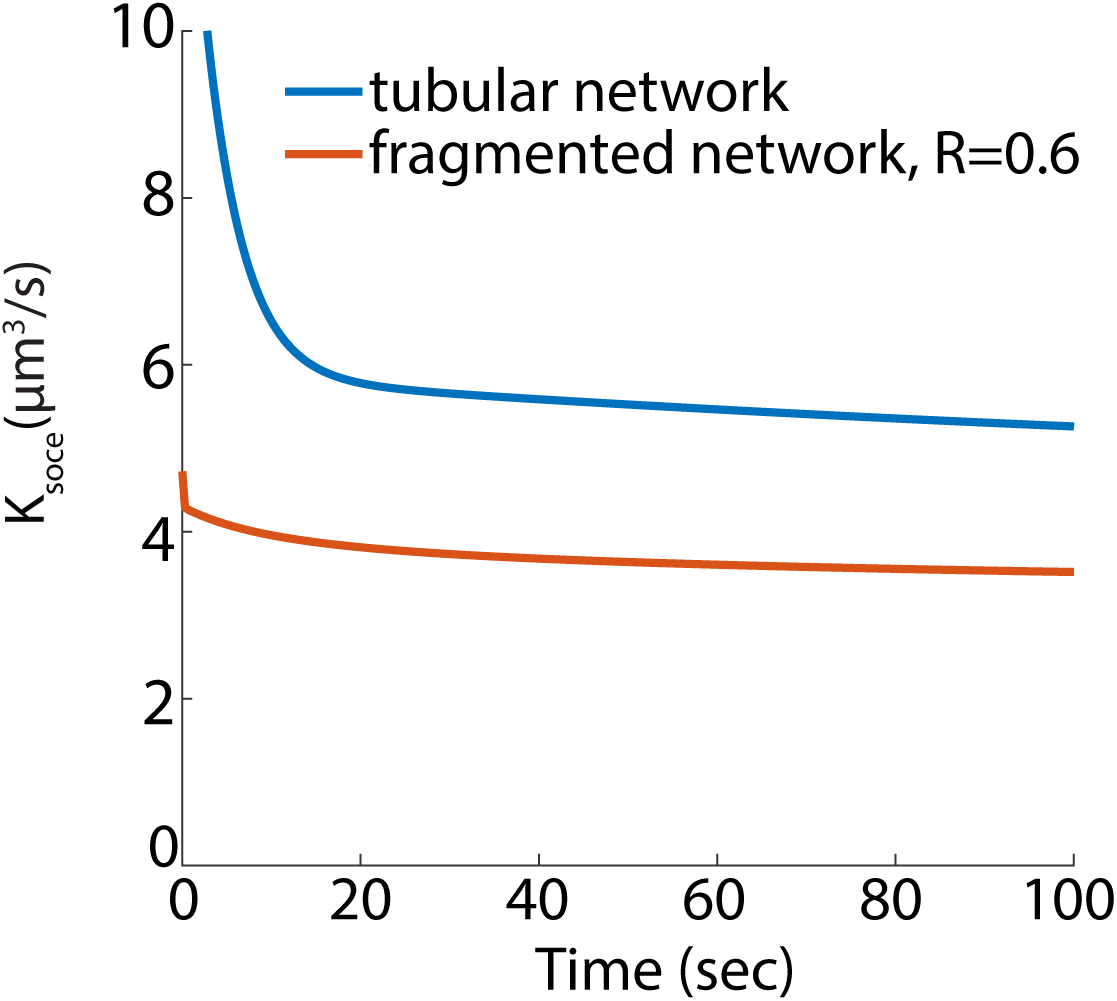
Modelled refill rates. Effective refill rate k_SOCE over time for a tubular network (WT) and a network of spheres with radius 0.6 µm (RTN3OE), extracted from 3D spatial simulations. Both networks have 40 peripheral contacts with the extracellular environment. The rate is initially high during peripheral ER filling and gradually plateaus as the peripheral network saturates and the reservoir continues to fill. We use the value of kSOCE at 10 seconds—6.5 µm³/s for WT and 4.0 µm³/s for RTN3OE—as input parameters for the aspatial model to reflect the quasi-steady-state refill rate.

**Figure S5:**
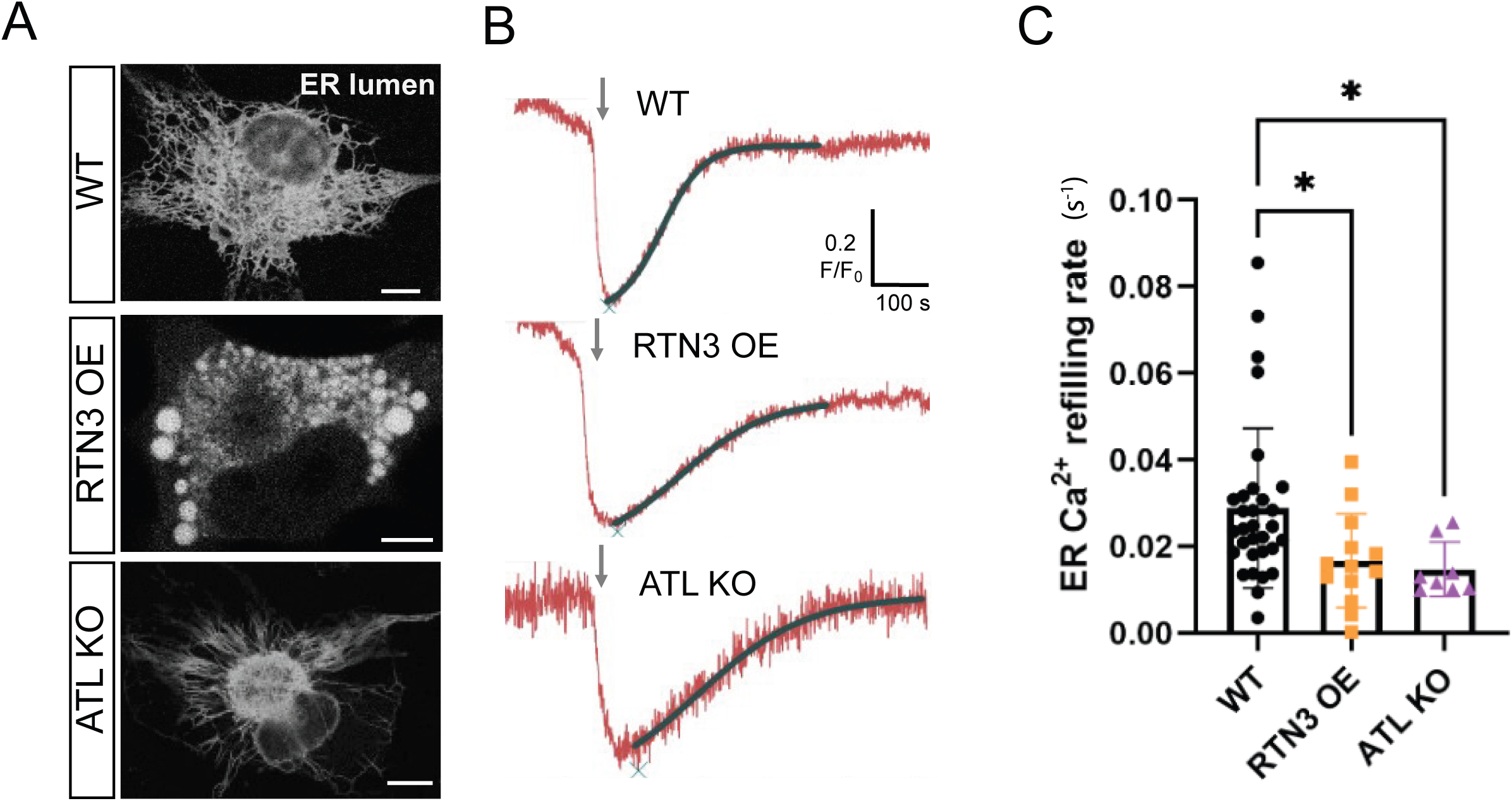
ER Ca^2+^ refill is slower in ATL KO ER networks. **A.** Micrographs of the ER lumen (visualised through R-CEPIA) in COS-7 cells WT (top), RTN3 OE (middle) and ATL KO (bottom). Scale bars: 10 µm. **B.** Normalized intensity profiles of an ER Ca^2+^ probe’s signal (ER-LAR-GECO1) over time in live COS-7 cells WT (top), RTN3 OE (middle) and ATL KO (bottom). Histamine (100 µM) addition is indicated by the arrow. Cytosolic Ca^2+^ was buffered with Bapta-AM (100 µM) to promote spatial globalisation of Ca^2+^ release. **C.** Quantification of ER Ca^2+^ refilling rates, obtained from fitting the recovery portion of the intensity profiles (sigmoid fits indicated by dark green lines in (B)). Each dot represents the refill rate for a single cell (WT: n= 31, RTN3 OE: n=13, ATL KO: n=8 cells. *p < 0.05, KW test).

Video 1. iNeurons’ spontaneous synchronous cytosolic Ca^2+^ bursts, related to Figure 1C. Time-lapse imaging of cytosolic Ca^2+^ (detected through Neuroburst) in iNeurons after 28 days of differentiation.

Video 2. iNeurons’ cytosolic Ca^2+^ bursts depend on ER Ca^2+^, related to Figure 1D. Time-lapse imaging of cytosolic Ca^2+^ in iNeurons as in Video1, before (left) and after CPA treatment (25 µM, middle), and after washout (right). During CPA treatment - causing ER Ca^2+^ depletion - the synchronous cytosolic bursts stop.

Video 3. ER fragmented through RTN3aOE can’t support the iNeurons’ cytosolic Ca^2+^ bursts, related to Figure 2E-G. Time-lapse imaging of cytosolic Ca^2+^ of iNeurons as in Video1 (middle), with sparse exogenous expression of RTN3a-Halo (labelled with Halo-ligand JFX650, magenta, left). iNeurons expressing RTN3-Halo don’t fire.

Video 4. ER fragmented through NSG392EOE can’t support the iNeurons’ cytosolic Ca^2+^ bursts, related to Figure 2E-G. Time-lapse imaging of cytosolic Ca^2+^ of iNeurons as in Video1 (middle), with sparse exogenous expression of Halo-NSG392E (labelled with Halo-ligand JFX650, magenta, left). iNeurons expressing Halo-NSG392E don’t fire.

Video 5. Loss of bursts synchrony after ER fragmentation, related to Figure 2H-J. Time-lapse imaging of cytosolic Ca^2+^ of iNeurons as in Video1, before (left) and after (middle) abundant exogenous expression of RTN3a-Halo (labelled with Halo-ligand JF646, magenta, right). After expressing RTN3a-Halo, the iNeurons’ network loses synchrony.

Video 6. ER fragmentation affects ER Ca^2+^ refill. Time-lapse imaging of ER Ca^2+^ in iNeurons detected through GCaMP6ER-150 (left) after emptying of ER Ca^2+^ through BTP2 and washing it out. The iNeuron expressing RTN3-Halo (labelled with Halo-ligand JF646, right) recover its ER Ca^2+^ slower than the one not expressing it. Timestamp is in format mins : secs. Scalebar: 20 µm.

Video 7. iNeurons’ cytosolic Ca^2+^ bursts depend on ER refill through SOCE, related to Figure 4A. Time-lapse imaging of cytosolic Ca^2+^ in iNeurons as in Video1, before and after BTP2 treatment (10 µM). After adding BTP2 – which inhibits SOCE - the synchronous cytosolic bursts gradually decrease in amplitude until completely stopping.

## Materials and Methods

**Table 1.**
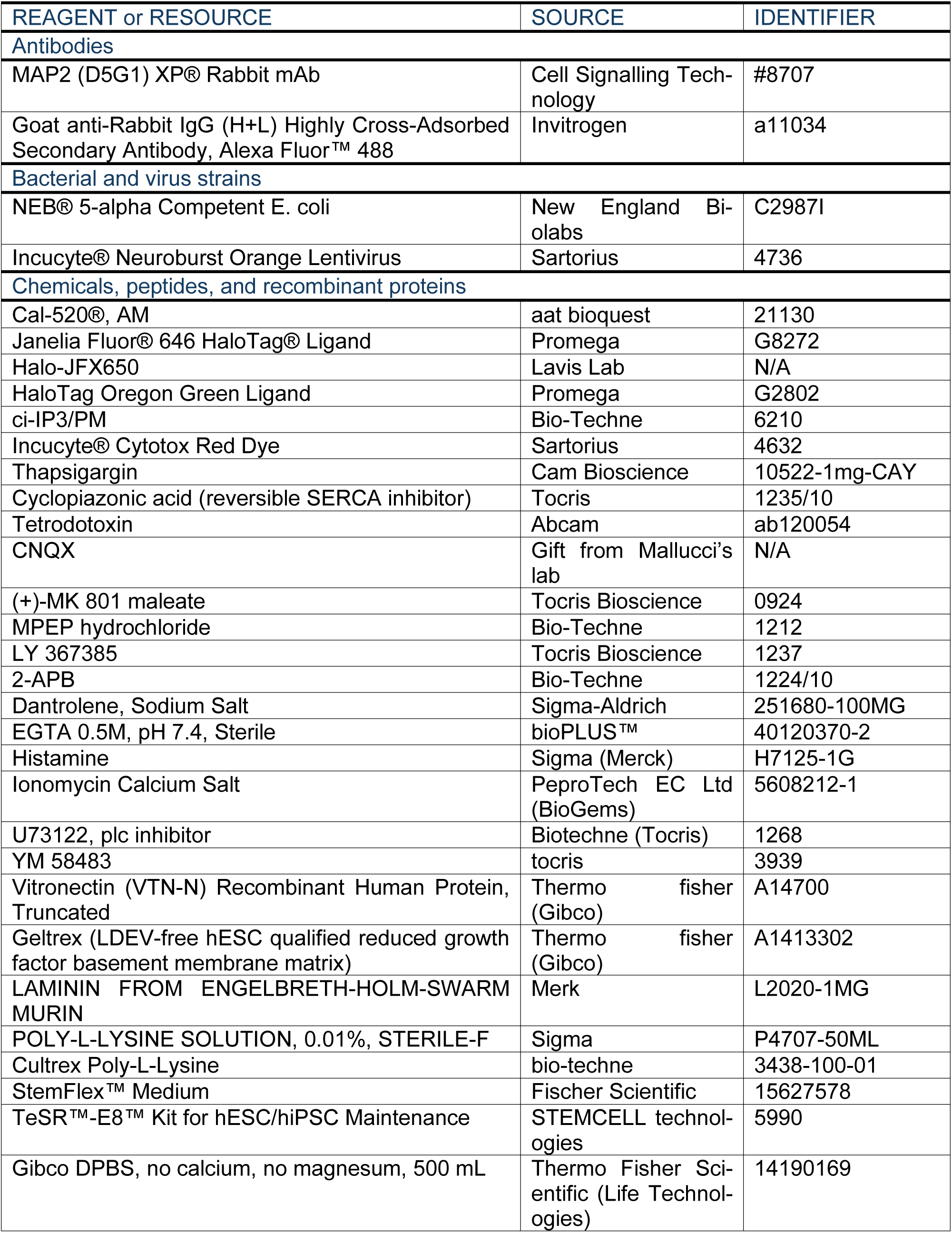
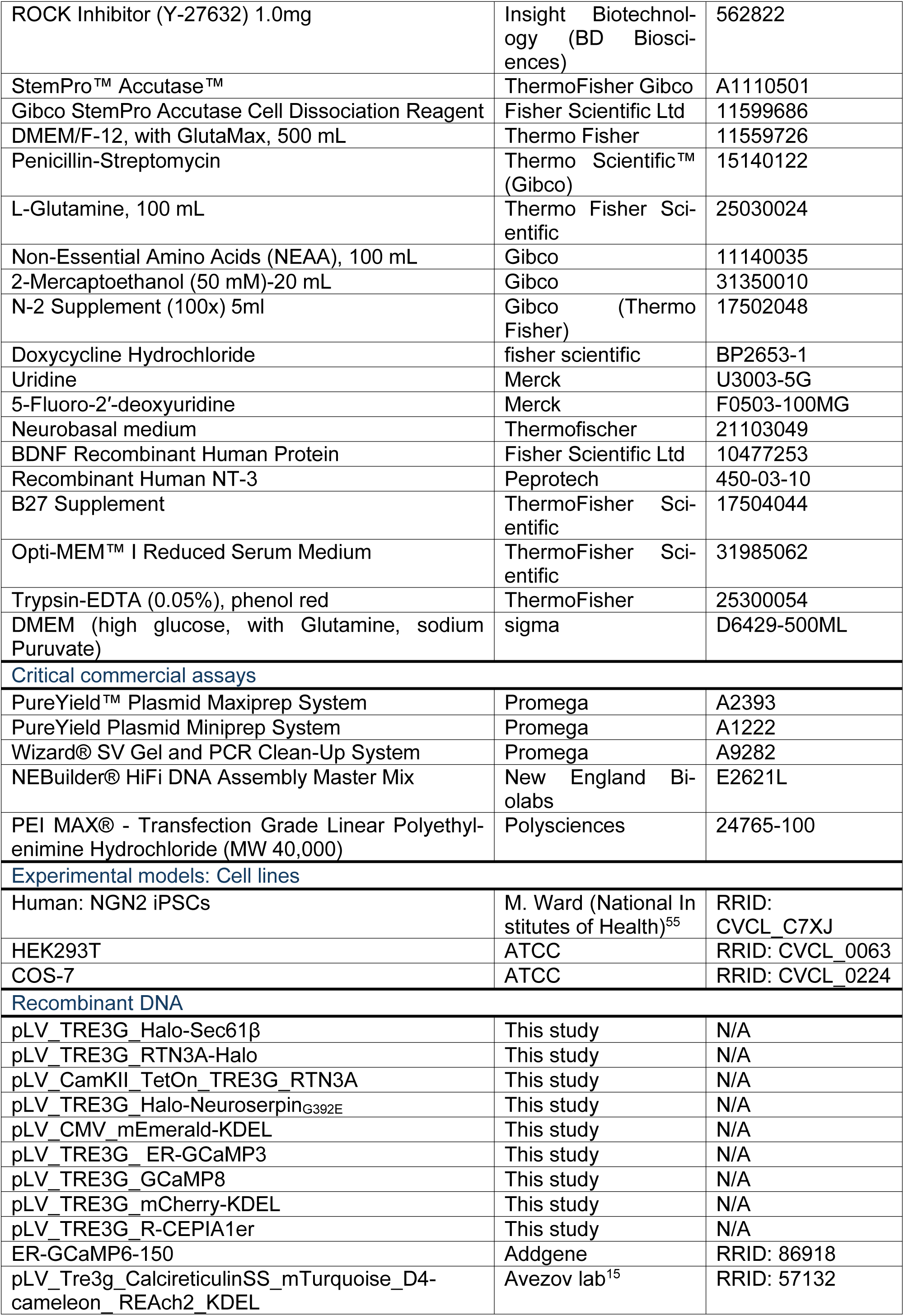
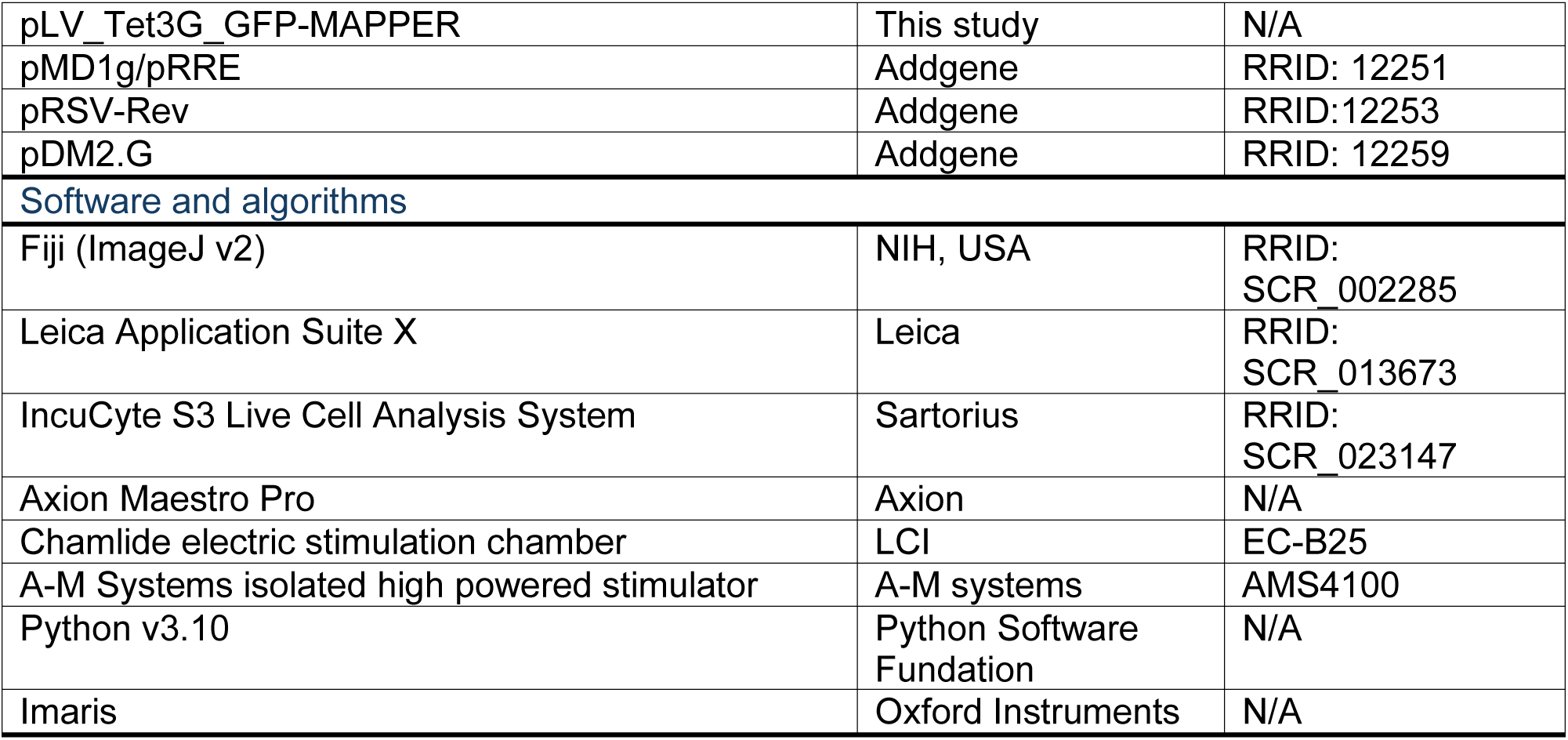
Key resources table.

### iPSCs maintenance and differentiation into iNeurons

Human Induced Pluripotent Stem Cells (hiPSCs) with stable integration of NGN2 into a safe-harbour locus under a doxycycline-inducible promoter (iNeurons)^55^, were cultured in STEM FLEX medium (STEMCELL technologies) on vitronectin (ThermoFisher Gibco)-coated plates (1:100 dilution, 1 hour) and treated with 10µM Rock Inhibitor (BD biosciences) in the first 24 hours after seeding. To induce neuronal differentiation, cells were transferred on Geltrex (ThermoFisher Gibco)-coated plates (1:100 dilution, 1 hour), and kept in STEM FLEX medium supplemented with Rock Inhibitor for 24 hours. Then, the STEM FLEX was replaced with DMEM/F12 (Gibco) supplemented with 10 µl/ml N-2 supplement (Gibco), 2mM L-glutamine (Gibco), 10 µl/ml NEAA (Gibco), 50 µM 2-mercaptoethanol, 10 µl/ml Pen-Strep (Gibco) and 1 µg/ml doxycycline (Fischer Scientific). This medium was changed daily for two days. On day 3 after induction, the medium was replaced with the maintenance medium Neurobasal (Gibco) supplemented with 20 µl/ml B-27 (Gibco), 2 mM L-glutamine (Gibco), 50 µM 2-mercaptoethanol (Gibco), 10 µl/ml penstrep (Gibco), 1ug/ml doxycycline (Fischer Scientific), 10 ng/ml NT-3 (Gibco) and 10 ng/ml BDNF (Gibco). At day 4, cells were transferred to different kinds of plates, depending on the analysis modality, coated with poly-L-lysine (Bio-Techne) overnight and laminin (Merk) at 1:100 dilution for 2 hours. The media was supplemented with 3 µM uridine (Sigma-Aldrich), 1 µM 5-Fluoro-2′-deoxyuridine (Sigma-Aldrich), and 10 µM ROCK inhibitor (BD biosciences) for 24 hours after seeding. The maintenance medium was subsequently changed every day for 4 days and every other day after day 8.

### MAP2 Immunostaining

Cells were fixed with a pre-warmed fixation buffer (2% PFA, 2% glutaraldehyde, 100 mM cacodylate (pH 7.4), and 2 mM CaCl_2_) for 30 min at room temperature (RT). After washing with 0.001% Triton in PBS, cell membrane was permeabilized with 0.5% Triton X-100 in PBS for 5 minutes. After, neurons were incubated in blocking solution (10% goat serum in PBS) for one hour. Then, after washing with PBS, samples were incubated overnight with primary antibody (1:500, MAP2 (D5G1) XP® Rabbit mAb, Cell Signalling Technology, #8707) in 10% goat serum at 4°C. On the next day, cell cultures were washed and incubated for one hour with secondary antibodies (1:1000 in PBS, Alexa 488, Invitrogen, goat anti-rabbit).

### Multielectrode array field potentials recordings

For multielectrode array (MEA) recordings, differentiated neurons were seeded on day 4 on a CytoView MEA 48 plate (Axion Biosystems, M768-tMEA-48W) coated as described above. 10,000 cells resuspended in 10 µl we added directly over the electrodes region in each well. Once attached, 300 µl of medium was added in each well and changed regularly as described above. Electrical activity was then recorded at ≥ day 21 using the Axis Navigator - Neuronal Activity Module of the Axion Maestro Pro (Axion Biosystems). Single electrode data was then extracted through the Axion Data Export tool and plotted using a custom Python code, while raster plots were generated through the Axion Neural Metric Tool.

### Whole-network cyto-Ca^2+^ recordings

For whole-network cytosolic Ca^2+^ recordings, differentiated neurons were seeded on day 4 on a 96-wells plate (Corning, 3595) at a density of 40,000 cells per well in 150 µl of maintenance medium. Ca^2+^ fluctuations of differentiated iNeurons were assessed using Neuroburst Orange, a lentivirus containing a genetically encoded Ca^2+^ indicator under a synapsin promoter (Sartorius, 4736). At day 7, neurons were transduced with 3 µl of Neuroburst lentiviral solution per well. Infected neurons were recorded at ≥ day 21 using the Neural Activity module of the automatic microscopy imaging system Incucyte (S3neuro, Sartorius). Wells were imaged at 3.33 Hz for 1 minute every 20 minutes using a 4× objective. The resulting videos were analysed using the Incucyte software, which calculated active objects counts and mean correlation values of each well. Means and SDs of these values per condition where then calculated and plotted using a custom Python code. The videos and masks of active objects were also extracted to plot the normalised Neuroburst intensity of each active object. Cells were treated by adding the drugs directly into the well, and washed-out by replacing medium 3 times. For the RTN3 OE experiment (Fig. 2G-I), cells were infected with RTN3 lentiviral particles at Day 8 and maintained in iN2 without DOX until the start of the experiment (day 28), when RTN3 OE was induced by adding DOX.

### Lentiviral particles production

8 x 10^6^ HEK293FT cells per flask were seeded in T175 flasks (Greiner) and incubated overnight. The next day, we incubated for each flask 2.5 µg of each third-generation viral packaging plasmids (Addgene #12251, #12253, #12259) and 2.5 µg of the plasmid of interest with 60 µl Polyethylenimine (PEI, Polysciences, 24765-100) in 1 ml Opti-MEM (Fisher Scientific) for half an hour and added the solution to the cell culture medium. The media (25 ml per flask) was then replaced every 24 h and collected after 48 and 72 hours. The collected media was filtered with a 0.45 μm filter and transferred into 50 ml tubes. 2 ml of 20% sucrose water solution was added at the bottom of each tube which were then centrifugated at 14000 rcf for 3h30min at 4°C. The formed pellet containing the viral particles was resuspended in 30 µl of PBS per 50 ml of collected media and stored at -80°C.

### Single-cell Cytosolic/ER Ca^2+^ recordings in firing iNeurons

For single-cell cytosolic cytosolic Ca^2+^ recordings, differentiated neurons were seeded on day 4 on a µ-Slide 8-well polymer bottom dish (IBIDI, 80801) coated as described above, at a density of 100,000 cells per well in 300 µl of maintenance medium. At day 15, wells were transduced with 6µl of Neuroburst (Sartorius, 6 µl per well) lentiviral particles. At day 28, cells were imaged on a STELARIS8 confocal microscope (Leica, Wetzlar, Germany) with a controlled environment (37°C, 5% CO_2_). ROIs were manually drawn around cells in the FOV in the Leica Stelaris software and their intensity values were extracted and plotted in Python.

For the EGTA experiment (Fig. 1H), videos were acquired at 2.3 Hz using a 10× objective, using a 558 nm excitation laser at 5% intensity, an emission detector set at 581-741 nm (50 gain) and the pinhole set at 11.31 AU (600 µm). During the recording (as indicated in the figures), cells were treated by adding EGTA (3 mM) directly into the well.

For the CPA experiment (Fig. 1I), ER Ca^2+^ fluctuations were imaged using ER-GCaMP6-150^56^, delivered through lentiviral transduction at day 4. Videos were acquired at 2.3 Hz using a 20× objective. The imaging setup was as follows: excitation lasers: 497 nm at 1% intensity, 552 nm at 2% intensity and the emission detectors were set at: 505-545 nm (50 gain) and 599-740 nm (50 gain). The pinhole was set to 5.62 AU (317.9 µm). Cells were treated during the recording (as indicated in the figures) by adding CPA (25 µM) directly into the well.

For recordings after ER-shape manipulation (Fig. 2E, F), media was replaced by iN2 without dox at day 5. At day 8, cells were transduced with lentivirus containing either RTN3a-Halo + mEmerald-KDEL (luminal marker to visualise ER vesicles) or Halo-NSG392E. To not affect neuronal maturation, the expression of the constructs under TRE3G promoter was induced through doxycycline addition (1 µg/ml) after checking that most of the cells in the well had Ca^2+^ bursts. After 1 week from overexpression induction, cells were stained with 1 µM Halo-JFX650 overnight and washed 3x with media, left to rest for 1h in the incubator, then washed 1x with medium. Videos acquired at 2.3 Hz using a 63× objective. The imaging setup was as follows: excitation lasers: 558 nm at 5 % intensity and 658 nm at 0.1 % intensity and the emission detectors were set at: 578 – 621 nm (50 gain) and 709 – 845 nm (2.5 gain). The pinhole was set to 2.88 AU (275.1 µm). The iNeurons cultures containing NSG392EOE cells were imaged at 2.3 Hz using a 20× objective. The imaging setup was as follows: excitation lasers: 558 nm at 4 % intensity and 658 nm at 0.5 % intensity and the emission detectors were set at: 578 – 621 nm (40 gain) and 691 – 845 nm (2.5 gain). The pinhole was set to 4.33 AU (413.2 µm). Intensity values for each cell in the FOV were extracted from the Leica Stelaris software and plotted in Python. Representative images were smoothed using Gaussian Blur in Fiji (ImageJ).

For the BTP2 experiment (Fig. 4A), videos were acquired at 2.7 Hz using a 40× objective. The imaging setup was as follows: excitation laser 558 nm at 1 % intensity and the emission detector was set at 585-838 (50 gain). The pinhole was set to 5 AU (386 µm). During the recording, cells were treated by adding BTP2 (10 µM) directly into the well.

### Imaging of ER shape manipulation

Differentiation of parental hiPSCs into iNeurons was induced as described above, and cells were seeded on day 4 on a µ-Slide 8-well polymer bottom dish (IBIDI, 80801) coated as described above, at a density of 50,000 cells per well in 300 µl of maintenance medium. On the same day, cells were transduced with lentivirus containing either RTN3a-Halo + D4ER-Tq (ER luminal marker) or Sec61b + mEmerald-KDEL (ER luminal marker) or Halo-NSG392E + ER-targeted GCaMP3 (ER membrane marker). After 1 week of overexpression, cells were stained with 200 nM Halo Ligand JF646 overnight and washed 3x with media, left to rest for 1 h in the incubator, then washed 1x with medium. Cells were imaged on a STELARIS8 confocal microscope (Leica, Wetzlar, Germany) with a controlled environment (37°C, 5% CO_2_) and imaged using a 63× objective. The imaging setup was as follows. For Sec61b + mEmerald: excitation lasers: 487 at 7% intensity and 646 nm at 7% intensity and the emission detectors were set at 498 nm – 618 nm (50 gain) and 659 nm – 834 nm (2.5 gain). The pinhole was set to 0.99 AU (95.1 µm). For RTN3a-Halo + D4ER-Tq: excitation lasers: 462 at 5% intensity and 646 nm at 5% intensity. The emission detectors were set at 472 – 558 nm and 661 – 834 nm. The pinhole was set to 0.99 AU (95.1 µm). For NSG392E OE + ER-targeted GCaMP3: excitation lasers: 488 nm at 5% intensity and 646 nm at 2% intensity. The emission detectors were set at 506- 560 nm and 658-776 nm. The pinhole was set to 0.6 AU (56.9 µm) and a line averaging of 2 was used.

### 3D reconstructions and ER-PM contacts quantification

To overcome the low transduction efficiency, MAPPER^48^ was transduced in parental hiPSCs, and MAPPER positive cells were selected through puromycin. Differentiation of the stable MAPPER overexpressing hiPSCs into iNeurons was induced as described above, and cells were seeded on day 4 on a µ-Slide 8-well polymer bottom dish (IBIDI, 80801) coated as described above, at a density of 50,000 cells per well in 300 µl of maintenance medium. On the same day, cells were transduced with lentiviruses containing mCherry-KDEL and either Halo- NSG392E or RTN3a-Halo. After 1 week of overexpression, cells were stained with 100 nM Halo Ligand JF646 overnight and washed 3x with media, left to rest for 1h in the incubator, then washed 1x with medium. Cells were then imaged on a STELARIS8 confocal microscope (Leica, Wetzlar, Germany) with a controlled environment (37°C, 5% CO_2_). Z-stacks were acquired using a 63× objective and Δz = 0.304 µm. The imaging setup was as follows: excitation lasers: 488 at 2% intensity, 487 at 1%, 647 nm at 1% intensity and the emission detectors were set at 496 - 547 nm, 601 - 634 nm and 682 – 850 nm. The pinhole was set to 0.99 AU (95.1 µm) and a line averaging of 2 was used. Representative 3D projections were generated through the LAS X 3D Visualization module. The raw z-stacks were analysed through the IMARIS software, where surfaces for MAPPER positive objects were created. The number of individual MAPPER clusters was then extracted as the number of resulting IMARIS surfaces and plotted in Python.

### Sample processing for electron microscopy and SEM imaging

Samples were fixed in fixative (2% glutaraldehyde/2% formaldehyde in 0.05 M sodium cacodylate buffer pH 7.4 containing 2 mM Ca^2+^ chloride) overnight at 4°C. After washing 5x with 0.05 M sodium cacodylate buffer pH 7.4, samples were osmicated (1% osmium tetroxide, 1.5% potassium ferricyanide, 0.05 M sodium cacodylate buffer pH 7.4) for 3 days at 4°C. After washing 5x in DIW (deionised water), samples were treated with 0.1% (w/v) thiocarbohydrazide/DIW for 20 minutes at room temperature in the dark. After washing 5x in DIW, samples were osmicated a second time for 1 hour at RT (2% osmium tetroxide/DIW). After washing 5x in DIW, samples were block stained with uranyl acetate (2% uranyl acetate in 0.05 M maleate buffer pH 5.5) for 3 days at 4°C. Samples were washed 5x in DIW and then dehydrated in a graded series of ethanol (50%/70%/95%/100%/100% dry), and 100% dry acetonitrile, 3x in each for at least 5 min. Samples were infiltrated with a 50/50 mixture of 100% dry acetonitrile/Quetol resin (without BDMA) overnight, followed by 3 days in 100% Quetol (without BDMA). Then, the sample was infiltrated for 5 days in 100% Quetol resin with BDMA, exchanging the resin each day. The Quetol resin mixture is: 12 g Quetol 651, 15.7 g NSA (nonenyl succinic anhydride), 5.7 g MNA (methyl nadic anhydride) and 0.5 g BDMA (benzyldimethylamine; all from TAAB). Samples were placed in embedding moulds and cured at 60°C for 2 days.

Thin-sections (∼ 100 nm) were cut using an ultramicrotome (Leica Ultracut E) and placed on melinex coverslips and allowed to air-dry. The coverslips were mounted on aluminium SEM stubs using conductive carbon tabs and the edges of the coverslips were painted with conductive silver paint. Then, samples were sputter coated with 30 nm carbon using a Quorum Q150 T E carbon coater.

Samples were imaged in a Verios 460 scanning electron microscope (FEI/Thermofisher) at 4 keV accelerating voltage and 0.2 nA probe current in backscatter mode using the concentric backscatter detector (CBS) in immersion mode at a working distance of 3.5-4 mm. Stitched maps were acquired using FEI MAPS software using the default stitching profile and 5% image overlap.

### Cytotoxicity assay

Differentiation of parental hiPSCs into iNeurons was induced as described above, and cells were seeded on day 4 on a 96 well plate (Corning, 3595) coated as described above, at a density of 20,000 cells per well in 150 µl of maintenance medium. At day 5, media was replaced by iN2 without dox. At day 8, cells were transduced with lentivirus containing either RTN3a-Halo, Halo-NSG392E, mEmerald-KDEL (as control for ER luminal protein overexpression) or Halo-Sec61b (as control for ER membrane protein overexpression). To not affect neuronal maturation, the expression of the constructs under TRE3G promoter was induced through doxycycline addition (1 µg/ml) at day 21. After 1 week from overexpression induction, cells were stained with 250 nM of Cytotox Red Dye (Sartorius) and Halo was labelled with 1 µM HaloTag Oregon Green Ligand for 1 hour and washed 3x with media, left to rest for 1h in the incubator, then washed 1x with medium. Cells were imaged using an Incucyte S3 (Sartorius) microscope using a 20x objective. The resulting images were analysed using the Incucyte software, which calculated the area of green + red objects and total green objects per well. The ratio between these two values was then calculated and plotted in Python.

### Free ER Ca^2+^ concentrations measurements through FLIM

Differentiation of parental hiPSCs into iNeurons was induced as described above, and cells were seeded on day 4 on a µ-Slide 8-well polymer bottom dish (IBIDI, 80801) coated as described above, at a density of 50,000 cells per well in 300 µl of maintenance medium. On the same day, iNeurons were transduced with the D4ER-Tq FRET-FLIM probe^15^, and either RTN3-Halo or Halo-NS_G392E_. After 1 week of overexpression, cells were stained with 100 nM JF646-HaloTag ligand overnight. iNeurons were imaged on a confocal microscope (SP8, Leica, Wetzlar, Germany) with a controlled environment (37°C, 5% CO2) and imaged with a 63x objective. The following parameters were applied: excitation/emission of 448 (20% power)/460-498 nm, pinhole: 0.5 AU (48 µm), scan speed: 400 Hz and settings were set to reach 5000 photons/pixel. Images were processed using Leica STELLARIS8 FLIM wizard. ROIs were manually drawn around individual cells and data fitted to a mono-exponential decay function. Lifetime values were converted to [Ca^2+^]_ER_ using the formula reported in the probe’s original paper^42^. The calibration of D4ER-Tq in iNeurons was performed by inducing maximum ER Ca^2+^ condition with addition of ionomycin (10 μM) and minimal ER Ca^2+^ condition was obtained by adding Thapsigargin (3 μM) to trigger full depletion of the Ca^2+^ stored in the ER.

### Total ER Ca^2+^ load measurements

Differentiation of parental hiPSCs into iNeurons was induced as described above, and cells were seeded on day 4 on a µ-Slide 8-well polymer bottom dish (IBIDI, 80801) coated as described above, at a density of 50,000 cells per well in 300 µl of maintenance medium. On the same day, iNeurons were transduced with RTN3a-Halo or Halo-NSG392E. After 1 week of overexpression, cells were stained with 100 nM JF646-HaloTag ligand overnight and then with 5 µM Cal-520 (AAT Bioquest) for 1 hour. iNeurons were imaged on a confocal microscope (SP8, Leica, Wetzlar, Germany) with a controlled environment (37°C, 5% CO_2_). Cells were imaged at 0.39 Hz using a 20× objective. The imaging setup was as follows. Excitation lasers: 499 nm at 2% intensity and 646 nm at 0.01% intensity. The emission detectors were set at: 506-617 nm (10 gain) and 653-795 nm (2.5 gain). The pinhole was set to 9.5 AU (600 μm). During the recordings, Thapsigargin (2 µM) was added directly into the well. ROIs were manually drawn around each cell in the FOV in the Leica Stelaris software and their intensity values were extracted and plotted against time in Python after smoothing with a Savitzky-Golay filter (order 1, window size 15, mode “mirror”) from Scipy. The area under the curve (AUC) was calculated using the “trapz” function from NumPy on the smoothed signal.

### ER Ca^2+^ release measurements through IP3 photo-uncaging

Differentiation of parental hiPSCs into iNeurons was induced as described above, and cells were seeded on day 4 on a µ-Slide 8-well polymer bottom dish (IBIDI, 80801) coated as described above, at a density of 50,000 cells per well in 300 µl of maintenance medium. On the same day, iNeurons were transduced with GCaMP8^57^ and RTN3a-Halo. After 1 week of over- expression, cells were stained with 100 nM JF646-HaloTag ligand overnight and then treated with 3 μM ci-IP3/PM (Tocris) for 3 hours. iNeurons were imaged on a confocal microscope (STELLARIS 8, Leica, Wetzlar, Germany) with a controlled environment (37°C, 5% CO_2_). Cells were imaged at 1.15 Hz using a 20× objective. The imaging setup was as follows. Excitation lasers: 480 nm at 3.5% intensity and 646 nm at 0.01% intensity. The emission detectors were set at: 489-552 nm (100 gain) and 689-834 nm (10 gain). The pinhole was set to 9.5 AU (600 μm). Following an acquisition of pre-photo-uncaging images (3 frames), photo-uncaging of caged-IP3 was achieved by illumination (405 nm, 100% laser power) of the whole field of view (775x775 μm) for a duration of 130 frames using the Fly mode in the FRAP wizard. ROIs were manually drawn around cells in the FOV in the Leica Stelaris software and GCaMP8 intensity values were extracted. Intensity at each frame was normalised by intensity before photo-uncaging and plotted against time in Python after smoothing with a Savitzky-Golay filter (order 1, window size 20, mode “mirror”) from Scipy. Peaks were detected using the “argmax” function from NumPy on the smoothed signal and the area under the curve (AUC) was calculated using the “trapz” function from NumPy on the smoothed signal.

### Luminal transport measurements through FRAP

Differentiation of parental hiPSCs into iNeurons was induced as described above, and cells were seeded on day 4 on a µ-Slide 8-well polymer bottom dish (IBIDI, 80801) coated as described above, at a density of 50,000 cells per well in 300 µl of maintenance medium. On the same day, iNeurons were infected with mCherry-KDEL and RTN3a-Halo or Halo-NS_G392E_. After 1 week of overexpression, cells were stained with 100 nM JF646-HaloTag ligand overnight. iNeurons were imaged on a confocal microscope (STELLARIS8, Leica, Wetzlar, Germany) with a controlled environment (37°C, 5% CO_2_). Cells were imaged using a 20× objective. For each recording, a circular region of radius ∼ 0.66 μm was selected in the peripheral region of a cell containing vesicular ER or normal ER network. The imaging setup was as follows: excitation/emission of 587 nm (0.2% power)/597-623, the pinhole was set to 47.8 μm (0.5 AU). A 646 nm laser was used to identify the transduced cells but not recorded during FRAP. The evolution of the fluorescence signal of mCherry-KDEL was recorded at 3.7 Hz. Following an acquisition of pre-photo-bleaching images (5 frames), photobleaching was performed inside the circular region for 2 frames (0.54 s) by setting the 587 nm laser power to 100%. The recordings were processed in ImageJ^58,59^ with the FRAP profiler v2 plugin (Hardin lab) with the following setup: mono-exponential fit, ROI1: bleaching ROI, ROI2: entire image. The resulting t-halves were plotted in Python.

### ER Ca^2+^ refill measurements

Differentiation of parental hiPSCs into iNeurons was induced as described above, and cells were seeded on day 4 on a µ-Slide 8-well polymer bottom dish (IBIDI, 80801) coated as described above, at a density of 50,000 cells per well in 300 µl of maintenance medium. On the same day, iNeurons were infected with ER-GCaMP6-150^56^ and RTN3a-Halo or Halo-NS_G392E_. After 1 week of overexpression, cells were stained with 100 nM JF646-HaloTag ligand overnight. iNeurons were imaged on a confocal microscope (STELLARIS8, Leica, Wetzlar, Germany) with a controlled environment (37°C, 5% CO_2_). The iNeurons cultures containing RTN3 OE cells were imaged at 1.13 Hz using a 40× objective. The imaging setup was as follows: excitation lasers: 496 nm at 1.5% intensity and 661 nm at 0.01% intensity and the emission detectors were set at: 510 – 554 nm (50 gain) and 713 - 834 nm (50 gain). The pinhole was set to 5 AU (386 µm). The iNeurons cultures containing NS_G392E_OE cells were imaged at 0.10 Hz using a 20× objective. The imaging setup was as follows: excitation lasers: 496 nm at 3% intensity and 646 nm at 0.01% intensity and the emission detectors were set at: 514-620 nm (50 gain) and 654-805nm (50 gain). The pinhole was set to 5.1 AU (288.9 µm). ROIs were manually drawn around cells in the FOV in the Leica Stelaris software and GCaMP6-150 intensity values were extracted. Values were plotted in Python after normalization 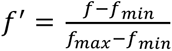. The sigmoid function 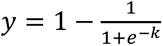 was fitted to this normalised trace using the Curve Fit function from Scipy, where k is the logistic growth rate and x_0_ is the x value of the function’s midpoint. X_0_ (time to half refill) was reported as a measure of refill speed.

### Cyto-Ca^2+^ recordings after refill inhibition through BTP2

Differentiation of parental hiPSCs into iNeurons was induced as described above, and cells were seeded on day 4 on a µ-Slide 8-well polymer bottom dish (IBIDI, 80801) coated as described above, at a density of 100,000 cells per well in 300 µl of maintenance medium. Cytosolic Ca^+^ fluctuations were imaged using Neuroburst Orange (Sartorius, 6 µl per well), delivered through lentiviral transduction at day 15. At day 28, cells were imaged on a STELARIS8 confocal microscope (Leica, Wetzlar, Germany) with a controlled environment (37°C, 5% CO_2_). Cells were imaged at 2.7 Hz using a 40× objective. The imaging setup was as follows: excitation laser 558 nm at 1% intensity and the emission detector was set at 585-838 (50 gain). The pinhole was set to 5 AU (386 µm). During the recording, cells were treated by adding BTP2 (10 µM) directly into the well. Intensity values for each cell in the FOV were extracted from the Leica Stelaris software and plotted in Python.

### Field stimulation

Differentiation of parental hiPSCs into iNeurons was induced as described above, and cells were seeded on day 4 on 25 mm glass coverslips coated as described above, at a density of 1x106 cells per well in 2 ml of maintenance medium. At day 5, media was replaced by iN2 without dox. At day 8, cells were transduced with RTN3a-Halo + mCherry-KDEL (luminal marker to visualize ER vesicles). To not affect neuronal maturation, the expression of the RTN3-Halo under TREG promoter was induced through doxycycline addition (1 µg/ml) after day 20. After 1 week from overexpression induction, iNeurons were stained with 5 μM Cal- 520 and 200 nM JF646 Halo-ligand for 1 hour. Coverslips were mounted into an EC-B25 stimulation chamber (Chamlide) and imaged using a Nikon Ti2 widefield microscope using a custom stage insert and 20x objective at 37°C and 5% carbon dioxide. Stimulation was performed using an AMS4100 stimulator (A-M Systems) and dedicated software. The field of view was adjusted to encompass cells with and without overexpression of RTN3. Repetitive stimulation experiments were conducted using a biphasic 1ms waveform and a voltage setting of 16 V. iNeurons were stimulated with bursts of 50 Hz stimulation, each lasting 0.2 seconds, at 15- second intervals.

### Cyto and ER Ca^2+^ recordings in COS7 cells during EGTA + Histamine Treatment

COS-7 cells stably expressing the Tet-On element were transduced with GCaMP8^57^ and RCEPIA_ER_^60^. Cells were then seeded on a 4 well dish (Greiner Bio-One, 627975) and expression of the Ca^2+^ sensors was induced by adding Dox (1 µg/ml) for 3 days. Cells were imaged on a STELARIS8 confocal microscope (Leica, Wetzlar, Germany) with a controlled environment (37°C, 5% CO_2_). Cells were imaged at 0.78 Hz using a 20× objective. The imaging setup was as follows: excitation lasers: 480 nm at 1% intensity, 562 nm at 2% intensity and the emission detectors were set at: 493-550 nm (50 gain) and 611-834 nm (50 gain). The pinhole was set to 10.47 AU (600 µm). Cells were pre-incubated with EGTA (3 mM) for 5 minutes and Histamine (100 µM) was added during the recording (as indicated in figure). ROIs were manually drawn around cells in the FOV in the Leica Stelaris software and intensity values were extracted and plotted in Python.

### ER Ca^2+^ refill in COS7 cells after Histamine-induced release

For global ER Ca^2+^ release induced by Histamine, cells were transfected by electroporation using the Neon Transfection System (Invitrogen) to express ER-LAR-GECO1 (together with RTN3E-Halo for RTN3 OE cell line). Cells were imaged 48-72 hours post-transfection on a confocal microscope (STELLARIS8, Leica, Wetzlar, Germany) with environmental control of the stage (37◦C, 5% CO2). Cells were imaged at 2 Hz using a 20× objective The imaging parameters were: Excitation / emission-GCaMP3: 480 / 485−550 nm, ER-LAR-GECO1: 550 / 600−670 nm, JF646: 646 / 670−750 nm. Cells were incubated with BAPTA-AM (100 μM, 15 minutes) and histamine (100 μM) was added during recording (as indicated in figure). Images were acquired for 5 minutes post-treatment. Image analysis was done using Fiji, and a custom code written in Python. Cells were manually segmented and fluorescence intensity time series within cell outlines were extracted using Fiji and normalised to the intensity measured at t_0_. ER Ca^2+^ drops were automatically detected in intensity traces using Python’s function find peak. Intensity traces were smoothed using a moving average with an 8- frame window. Refill rates were obtained by fitting a sigmoid curve to the smoothed intensity traces, from the lowest value to an end point defined by the length of the fit parameter during analysis: 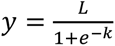, where L is the amplitude of the curve fit and k the logistic growth rate, with k reported as the refill rate.

### Statistical tests

All statistics were computed in Python using the ttest_ind function from Scipy. All values are reported as average +/- STD except when indicated otherwise.

### Modelling

#### Transport through network of spheres

We estimate the rate with which diffusive particles move between two spheres of volume *V* connected by a tube of radius *r* and length *l* by finding the mean first passage time *τ* for a particle starting uniformly distributed in one sphere to enter the second sphere. In the limit where *r* ≪ *l* ≪ *R*, this time has been previously worked out as^61^

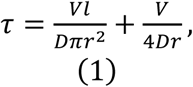

where *V* is the sphere volume. This result can also be derived through an electrostatic analogy^62^ by adding in series the diffusive resistance for escaping through a narrow pore and for transitioning across a straight tube. The timescale to refill an empty sphere connected through tubules to *n* neighbouring spheres is simply *π*/*n*.

For a well-connected tubular lattice, the effective three-dimensional diffusivity of particles embedded in the network is given by 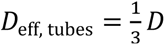, where *D* is the diffusivity inside the lumen.

The factor of 1*/*3 arises because the particles are only able to move along one dimension at a time when diffusing through the tubules.

For a lattice of vesicles with radius *R* connected by tubules of length *l*, we can approximate the particle motion as a series of discrete hops (transitions between spheres), each requiring time *τ*/*n* and each shifting the particle by distance 2*R* + *l* along a single dimension. The resulting effective diffusivity is then

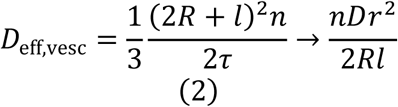

in the limit of *r* ≪ *l* ≪ *R*. For the vesicular lattice considered here (with *r* = 0.018 µm, *R* = 0.6 µm, *l* = 0.3 µm, *n* = 6), the ratio of effective diffusivities is *D*_eff, vesc_/*D*_eff, tubes_ ≈ 0.016. This estimate implies that diffusive particles will move much more slowly across an unbounded network of vesicles connected by narrow tubules (as for the RTN3OE ER structure) than they will in a purely tubular network (as in the WT ER structure). The vesicles serve as traps for diffusive particles, which must find narrow tubule entrances to escape. Larger spheres are expected to yield slower transport, as observed experimentally for the comparison of RTN3OE and NS architectures.

#### Simulations of network Ca^2+^ refill

To specifically explore the effect of Ca^2+^ transport and network structure on the refill process, we made use of a spatially resolved model that eschews all other aspects of Ca^2+^ dynamics. This model treats the WT ER as a 3D network of narrow tubules, enclosed within a spherical cell, with a finite number of contacts to the plasma membrane at the cell periphery. At these contacts, SOCE is assumed to be rapid enough to maintain a fixed free Ca^2+^ concentration of 0.5mM within the ER lumen. Diffusive transport of both free and bound Ca^2+^ ions then determines the typical rate of refilling for the entire network. The default number of PM contact sites (in Fig. 3G, H) is set to 40.

##### Network construction

The 3D tubular network is constructed by starting with a diamond lattice and randomly shifting one edge at each node to leave junctions of degree 3. The average edge length is 1 µm. The network is cropped to lie within a spherical shell of outer radius 10 µm (representing the cell boundary) and inner radius 5 µm (representing the nucleus). The network is then connected to a well-mixed perinuclear reservoir with a volume of 189 µm^3^, corresponding to 10 sheets of thickness 0.06 µm surrounding a nucleus of radius 5 µm.

For the partially fragmented ER structure, we begin with a hexagonally close packed lattice of nodes and randomly disconnect edges to reach an average degree 6 at each node. The individual nodes then represent well-mixed spherical vesicles. We set the vesicle radius to 0.6 µm for the RTN3OE structure and to 0.5 µm for the NS structure, based on measurements of the ER bubbles observed in 2D images (as in Fig. 2B). Each bubble was connected to its (on average) 6 neighbours by a tubule of length 0.3 µm. The tubule radius was selected based on the observed rate of fluorescent protein recovery following photobleaching of an individual bubble (Fig. 3C). Using the analytic approximation for the rate of transfer between individual spheres (Eq. 1), and assuming a connectivity of *n* = 6, tubules of radius 0.018 µm results in an average refill time of *τ*/*n* = 16 sec and *τ*/*n* = 9 sec for spheres of radius 0.6, 0.5 µm, respectively. As for the purely tubular case, the vesicular network was connected by short tubules to a perinuclear reservoir.

##### Buffer binding

In both WT and fragmented networks, the intra-luminal space is assumed to contain buffer sites with binding strength *K_D_* = 0.2 mM. For the WT network, the total concentration of buffer sites is set to *S* = 2.7 mM, such that the total Ca^2+^ concentration in the filled lumen is five-fold higher than the free Ca^2+^. While the vesicular network has a larger total volume than the tubular one, we assume that the total number of buffer proteins remains the same across both morphologies. Consequently, the buffer concentration was scaled down in proportion to the volume ratio between the two.

Buffer proteins were taken to have a diffusivity of *D_b_* = 3 µm^2^*/*s^16^, while free Ca^2+^ was taken to diffuse ten-fold faster at *D_c_* = 30 *µ*m^2^*/*s. The concentration field of free Ca^2+^ is propagated forward using a finite volume approach, under an assumption of rapidly equilibrated binding, as described in prior work^15^.

##### Extracting refill rate estimate

To estimate a single rate constant for refill from the simulations, we track both ⟨*U*(*t*)⟩ (the free Ca^2+^ concentration averaged across the entire ER lumen) and the current *I*(*t*) of Ca^2+^ ions entering through the PM contact sites. The effective refill rate constant is then computed as *k*_SOCE_ = *I*(*t*)*/*(*c*_cs_ − ⟨*U*(*t*)⟩), where *c*_cs_ is the fixed free Ca^2+^ concentration at the PM contacts. Following an initial rapid peripheral filling phase, *k*_SOCE_(*t*) reached a plateau during a quasi-steady-state where peripheral Ca^2+^ became saturated, but the reservoir remained only partially filled.

#### Mathematical model for Ca^2+^ oscillations

To describe the dynamics of periodic Ca^2+^ pulses, we implemented an aspatial model that encompasses the interchange of ions between two well-mixed pools: the ER lumen and the cytoplasm. Our approach modifies a classic simplified model developed by Li and Rinzel^50^, which incorporates dynamic Ca^2+^-dependent activation and inactivation of IP3 receptors, as well as SERCA pumping and clearance from the cytoplasm. This model is extended here to explicitly include Ca^2+^ buffers in the ER lumen and refilling of the ER through SOCE.

The model tracks the dynamics of *u*_ER_ (concentration of free Ca^2+^ in the ER lumen) and *c*_cyto_ (cytoplasmic Ca^2+^ concentration). The total luminal Ca^2+^ is set to *c*_ER_ = *u*_ER_[1+*S/*(*u*_ER_ +*K_D_*)]. Cytoplasmic buffering is not included. The concentrations evolve based on Ca^2+^ fluxes according to

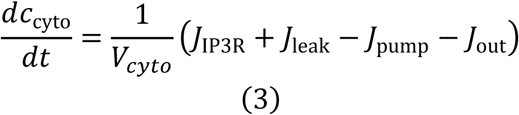

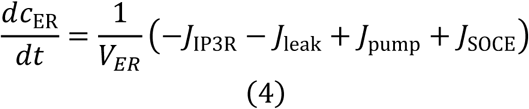

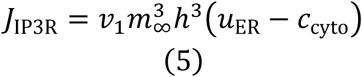

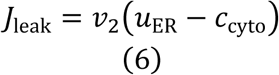

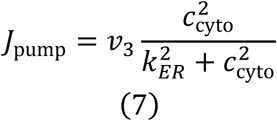

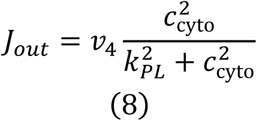

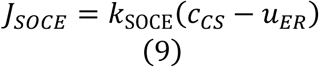

*J*_IP3R_ defines the flux through IP3-gated Ca^2+^-responsive Ca^2+^ channels. Each IP3 receptor subunit is assumed to contain one regulatory site for IP3 binding, one for Ca^2+^ activation, and one for Ca^2+^ inhibition. The probability of the channel being open depends on the state of each of the regulatory sites. The IP3 binding site and the Ca^2+^ activation site are assumed to equilibrate instantaneously to the steady state level:

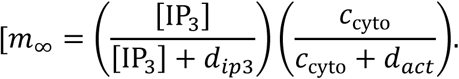

The inhibitory site responds to Ca^2+^ more slowly, and its state is defined by the Hodgkin-Huxley-like gating variable *h*(*t*), which evolves according to

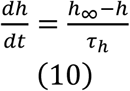

with steady-state h_inf and time constant tau:

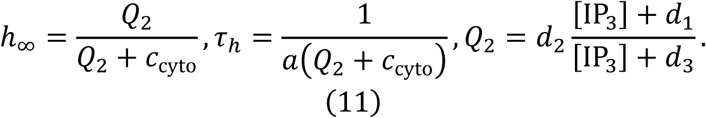

The cubic dependence of the regulatory sites was previously extracted from experimental fits^50^.

The flux *J*_leak_ accounts for passive leakage of Ca^2+^ from the ER to the cytoplasm, set to 0 in our version of the model. *J*_pump_ denotes active reuptake of Ca^2+^ into the ER by the SERCA pump, which has two active Ca^2+^-binding sites. *J*_out_ describes Ca^2+^ clearance from the cytoplasm through the plasma membrane via PMCA pumps that also have two binding sites.

In addition to these terms in the original Li & Rinzel model, we include the flux *J*_SOCE_, which describes the store-operated Ca^2+^ entry pathway, filling the ER lumen with Ca^2+^ brought in at plasma membrane contacts. We do not resolve the detailed dynamics at the plasma membrane contacts but rather assume that SOCE entry operates in a rapid and regulated fashion to maintain a fixed concentration of free luminal Ca^2+^ at the contact sites (*c_cs_*). In this simplified model, we take the filling of the bulk ER lumen to proceed at a rate proportional to the difference between this fixed contact site concentration and the average luminal free Ca^2+^. The rate constant *k*_SOCE_ is extracted from our spatially resolved simulations, using the value at time *t* = 10sec after refill is initiated. We note that this refill rate is already close to the long-time plateau value (see Fig. S3B4).

The rate of evolution for the free ER luminal Ca^2+^ is then found via the rapid-binding assumption:

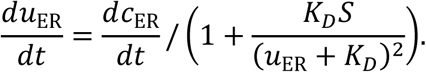

The half-max binding concentrations for all regulatory sites on IP3R channels have been previously estimated, and we use the published values (see Table 2). The buffer binding strength *K_d_* and total capacity *S* has also been estimated from experimental data^15^. We fix the free Ca^2+^level at contact sites based on an approximate measurement of ER free luminal Ca^2+^ in nonfiring cells (measured at 0.177 mM in WT iNeurons, Supplemental Fig. S2E). The refill rate constant *k*_SOCE_ is determined from spatially resolved simulations, and the ER and cytoplasmic volumes *V*_ER_*, V*_cyto_ are taken from our constructed WT network structures. Because we seek to isolate the effect of refill rate on Ca^2+^ oscillations, we use the same value of *V*_ER_ for both types of networks and alter only the refill rate constant. The remaining parameters: *v*_1_*, v*_3_*, v*_4_*, k*_ER_*, k*_PL_*, a* are tuned to demonstrate the plausibility of refill-rate-dependent oscillations as shown in Fig. 4.

##### Implementation

The aspatial model described above was implemented in Matlab, with the variables *u*_ER_(*t*)*, c*_cyto_(*t*)*, h*(*t*) evolved forward in time using the built-in ode45 function. Code for running this model is provided at at https://github.com/lenafabr/aspatialCaERrefill.

| Symbol | Meaning | Value | Rationale |
| --- | --- | --- | --- |
| $d_{ip3}$ | IP3R IP3 site $K_d$ (no inhibition) | 0.13 $\mu\text{M}$ | 63 |
| $d_2$ | IP3R $\text{Ca}^{2+}$ inhibition site $K_d$ | 1.049 $\mu\text{M}$ | 63 |
| $d_3$ | IP3R $\text{IP}_3$ site $K_d$ (with inhibition) | 943.4 nM | 63 |
| $d_{act}$ | IP3R $\text{Ca}^{2+}$ activation site $K_d$ | 82.34 nM | 63 |
| $k_D$ | $\text{Ca}^{2+}$-buffer protein dissociation constant | 0.2 $\mu\text{M}$ | 15 |
| $S$ | Buffer protein site conc. | 2.7 mM | 15 |
| $c_{cs}$ | $\text{Ca}^{2+}$ at SOCE contact site | 200 $\mu\text{M}$ | current measurements |
| $V_{ER}$ | ER volume | 212 $\mu\text{m}^3$ | model WT tubular network |
| $k_{SOCE}$ (WT) | SOCE refill rate constant (WT) | 6.5 $\mu\text{m}^3/\text{s}$ | spatial sims |
| $k_{SOCE}$ (RTN3OE) | SOCE refill rate constant (RTN3) | 4 $\mu\text{m}^3/\text{s}$ | spatial sims |
| [IP3] | IP3 concentration | 0.4 $\mu\text{M}$ | 50 |
| $v_1$ | Max $\text{IP}_3\text{R}$ flux rate | 69 $\mu\text{m}^3/\text{s}$ | Tuned for amplitude |
| $v_2$ | Passive $\text{Ca}^{2+}$ leak rate | 0 $\mu\text{m}^3/\text{s}$ | Tuned |
| $v_3$ | SERCA uptake max rate | 496 $\mu\text{M} \cdot \mu\text{m}^3/\text{s}$ | Tuned |
| $v_4$ | Cytoplasmic clearance rate | 368 $\mu\text{M} \cdot \mu\text{m}^3/\text{s}$ | Tuned |
| $k_{ER}$ | SERCA half-max $\text{Ca}^{2+}$ | 0.117 $\mu\text{M}$ | Tuned |
| $k_{PL}$ | Clearance half-max $\text{Ca}^{2+}$ | 0.106 $\mu\text{M}$ | Tuned |
| $a$ | IP3R $\text{Ca}^{2+}$ inhibition rate const. | 0.594 ($\mu\text{M} \cdot \text{s})^{-1}$ | Tuned |

## Acknowledgments

This work is supported by the UK Dementia Research Institute (award UK DRI-2004) through UK DRI Ltd., principally funded by the Medical Research Council (to E.A.). E.A. was further supported by Evelyn Trust and Alzheimer’s Society grant AS-525 (AS-PhD-19a-015). V.D. was supported by the Hughes Hall Cambridge Edwin Leong Scholarship in Life Sciences, Cambridge Trust and the Chui Wen Mei Fund at Hughes Hall. E.F.K. was supported by NSF grant #2034482.

